# Metabolic homeostasis and growth in abiotic cells

**DOI:** 10.1101/2022.10.16.512448

**Authors:** Amir Akbari, Bernhard O. Palsson

## Abstract

Metabolism constitutes the core chemistry of life. How it began on the early Earth and whether it had a cellular origin is still uncertain. A leading hypothesis for life’s origins postulates that metabolism arose from geochemical CO_2_-fixing pathways, driven by inorganic catalysts and energy sources, long before enzymes or genes existed. The acetyl-CoA pathway and the reductive tricarboxylic acid cycle are considered ancient reaction networks that hold relics of early carbon-fixing pathways. Although transition metals can promote many steps of these pathways, whether they form a functional metabolic network in abiotic cells has not been shown. Here, we formulate a nonenzymatic carbon-fixing network from these pathways and determine its functional feasibility in abiotic cells by imposing the fundamental physico-chemical constraints of the early Earth. Using first principles, we show that abiotic cells could have sustainable steady carbon-fixing cycles that perform a systemic function over a relatively narrow range of conditions. Furthermore, we find that in all feasible steady states, the operation of the cycle elevates the osmotic pressure, leading to volume expansion. These results suggest that achieving homeostatic metabolic states under prebiotic conditions was possible, but challenging, and volume growth was a fundamental property of early metabolism.

---

How life originated on a lifeless planet from nothing but a few inorganic precursors is a fundamental question in biology. Ever since Oparin’s primordial soup theory [1], two major hypotheses have been developed to address this question: one centered around information and the other around chemical self-organization. Which was more fundamental to the emergence of life is still debated [2]; that metabolism and self-organization play a prominent role in extant biology is not. Interestingly, phylogenetic studies of all domains of life trace biochemistry to an ancient autotrophic network at the core of metabolism [3]. These studies point to a core network that would have been anaerobic [3], phosphate-free [4], and reliant on simple carbon and energy sources [3].

Recent theories of life’s origins posit that metabolism originated from a geochemical protometabolism that predated enzymes and genes [2, 5]. They consider early metabolic pathways that were thermodynamically favorable and promoted by naturally occurring catalysts (*e*.*g*., transition metals) and reducing compounds (*e*.*g*., H_2_, FeS) [6]. Alkaline hydrothermal vents (AHVs) in the Hadean ocean are believed to have harbored these early pathways, providing inorganic catalysts, simple carbon sources, and a continuous supply of energy and reducing power [5] (Fig. 1A). The steep redox and pH gradients in AHVs would have been sufficient to drive energy and carbon metabolism [7]. Experimental evidence suggests that transition metals and inorganic reducing agents can promote several core metabolic pathways [8–13], corroborating AHV theories. However, whether a self-sustaining network comprising a series of core metabolic reactions could have spontaneously arisen and operated nonenzymatically under hydrothermal-vent conditions is still unclear.

**Figure 1:**
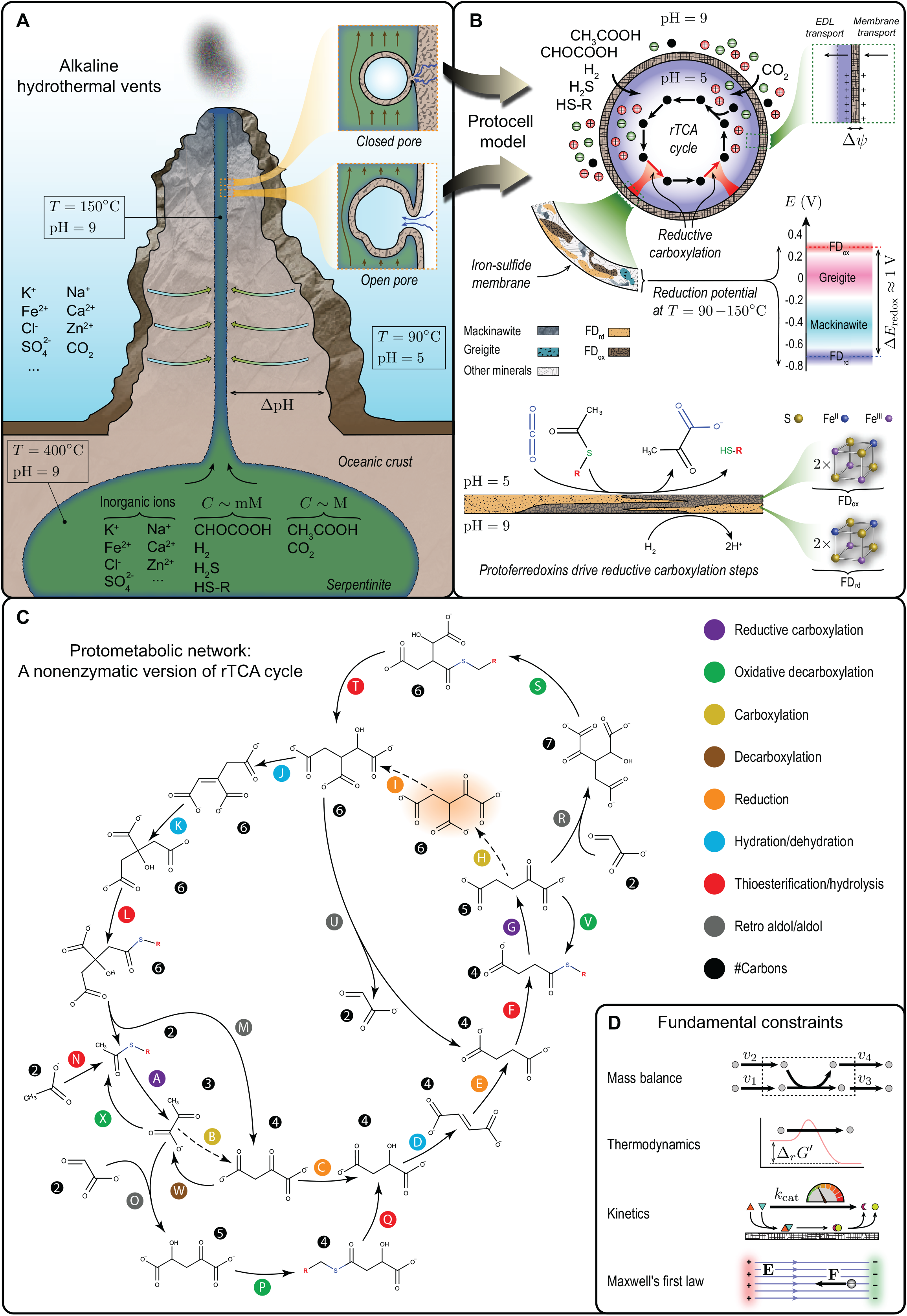
Emergence of first metabolic cycles from abiotic geochemical processes. (A) Deep-sea alkaline hydrothermal vents in the Hadean ocean were ideal environments from which metabolic pathways could spontaneously arise [5, 29]. Hydrothermal fluids formed by serpentinization would have been alkaline and rich in naturally occurring catalysts (*e*.*g*., metal ions, minerals) and reducing compounds (*e*.*g*., H_2_, H_2_S, FeS), providing the necessary ingredients for the emergence of metabolic networks. Thin-walled micropores made of iron sulfides forming along vent conduits, such as those shown in the inset (see Fig. S2), provide an interface between alkaline hydrothermal fluids and acidic ocean [7]. Redox and pH gradients across such inorganic barriers could have powered the synthesis of first organic molecules. (B) Protocell model with an iron-sulfide membrane simulates the formation of first metabolic cycles in vent micropores. All metabolic reactions beside carbon-fixing steps are catalyzed by transition metals in aqueous or solid phase dispersed inside the protocell in an acidic environment. Carbon-fixation reactions are catalyzed by protoferredoxins on the inner surface of the membrane. Protoferredoxins could have formed in the presence of iron and sulfur ions under hydrothermal conditions [7, 28]. Reduced (FD_rd_) and oxidized (FD_ox_) protoferredoxins would have had similar crystal structures to hydrothermal mineral redox couples (*e*.*g*., mackinawite/greigite), providing a sufficiently large redox potential to drive carbon fixation [7]. Reduced protoferredoxins consumed by carbon-fixation steps are regenerated on the outer surface of the membrane in an alkaline environment using H_2_ as a reducing agent. (C) Phosphate-free protometabolic network examined in this work comprising a nonenzymatic version of the rTCA cycle, containing all its intermediates except oxalosuccinate (highlighted in red). Dashed arrows indicate the corresponding enzymatic steps that are not included in the network. Thioesterification steps are driven by a simple hydrothermal thiol HS–R that could have been synthesized in sulphide-rich environments [7] with R a hypothetical substituent. (D) Fundumental constraints of the protocell model (see Supplementary Information for details).

At the origin of metabolism, boundary structures may have been required to (i) generate concentration gradients between compartments and their surroundings, providing a continuous supply of energy and material to maintain a stable far-from-equilibrium state [14] and (ii) create a barrier to limit diffusive loss of metabolic products [15]. Before the advent of enzymes, abiotic cellular structures could have facilitated the inception of first self-amplifying metabolic networks on Earth [16, 17]. However, the lack of selective transporters, efficient catalysts, and evolutionarily optimized energy-coupling means in such compartments could also have restricted the continued operation of these networks. Therefore, whether abiotic cells can nonenzymatically sustain stable carbon-fixing cycles is to be demonstrated. In this regard, first-principle analyses of abiotic systems that are consistent with established chemistries of nonenzymatic metabolism could elucidate the key factors constraining the emergence of metabolism.

Biology is a science of emergence. Therefore, we must show that if all components of abiotic cells work together collectively, a functioning system capable of reaching steady states (abiotic equivalents of homeostatic states) will emerge. Thus, we developed a protocell model (Fig. 1B) to simulate AHV conditions on the early Earth and answer whether AHV micropores could have harbored nonenzymatic autotrophic pathways using first principles (see Supplementary Information). In this protocell, the membrane is made of iron sulfides, a fraction of which are protoferredoxins. The membrane separates an acidic interior from alkaline surroundings. This pH gradient induces a positive membrane potential. It alleviates the dissipation of metabolic products [18] and provides the energy required for carbon fixation—a primitive energy-coupling mechanism mediated by protoferredoxins independently of ATPase and proton pumps. Abiotic metabolic reactions occur inside the protocell, and compounds cross the membrane unselectively *via* passive diffusion.

We began by assessing candidate abiotic chemical networks at the origin of metabolism, focusing on carbon assimilation in modern autotrophs. From the six known CO_2_-fixing pathways [2], the reductive tricarboxylic acid (rTCA) cycle is considered the most ancient autocatalytic and a primordial core network [19]. It contains the five universal precursors of metabolism (Fig. S3); all its intermediates but cofactors are phosphate-free; and several of its reactions can be promoted by transition metals [11]. Thus, we constructed a protometabolic network (Fig. 1C) based on the rTCA cycle to represent the first carbon-fixing cycles from which metabolism originated (see Supplementary Information). This network is cyclic and phosphate-free, only requiring inorganic cofactors and catalysts. Except for succinate reductive carboxylation, all its steps are derived from experimentally demonstrated nonenzymatic reactions or their close analogues [9, 11, 12, 20–22]. Neutral carboxylation steps from the enzymatic rTCA cycle (Fig. 1C, steps B and H) are not included in the network. These steps have not been reproduced nonenzymatically and are unlikely to proceed without evolutionarily tuned cofactors like ATP. Hence, they are bypassed through aldol and decarboxylation reactions. Acetate, glyoxylate, and CO_2_ are the carbon sources: Glyoxylate drives the foregoing bypass pathways, while acetate amplifies the cycle.

Were there plausible prebiotic conditions under which carbon fixation was feasible subject to the restrictions of enzyme-free metabolism and fundamental physico-chemical constraints? Could abiotic cells have operated on simple carbon and energy sources, while exhibiting fundamental characteristics of living cells, such as growth and homeostasis? Answering these questions is a challenging task, requiring a systems-level analysis of constrained reaction-diffusion problems. To addressed these questions, we formulated a dynamic system, accounting for the fundamental constraints [18, 23] arsing from mass conservation, thermodynamics, kinetics, and Maxwell’s first law (Fig. 1D). By studying the steady-state solutions and stability of this system, we quantified the feasibility and homeostatic states of the proposed protometabolism.

The turnover number and membrane potential are key parameters that encapsulate the restrictions of nonenzymatic metabolism. Low turnover numbers characterize inorganic-catalyst efficiencies, while positive membrane potentials are required to lower metabolic-product dissipation through semi-permeable inorganic membranes. Remarkably, we identified a restricted region in this parameter space where stable steady states that satisfy the fundamental constraints of the early Earth can be achieved (Fig. 2A). These states represent abiotic metabolic homeostasis, where a complete-cycle flux can be sustained. The stability region is bounded at low and high membrane potentials but unbounded in turnover numbers.

**Figure 2:**
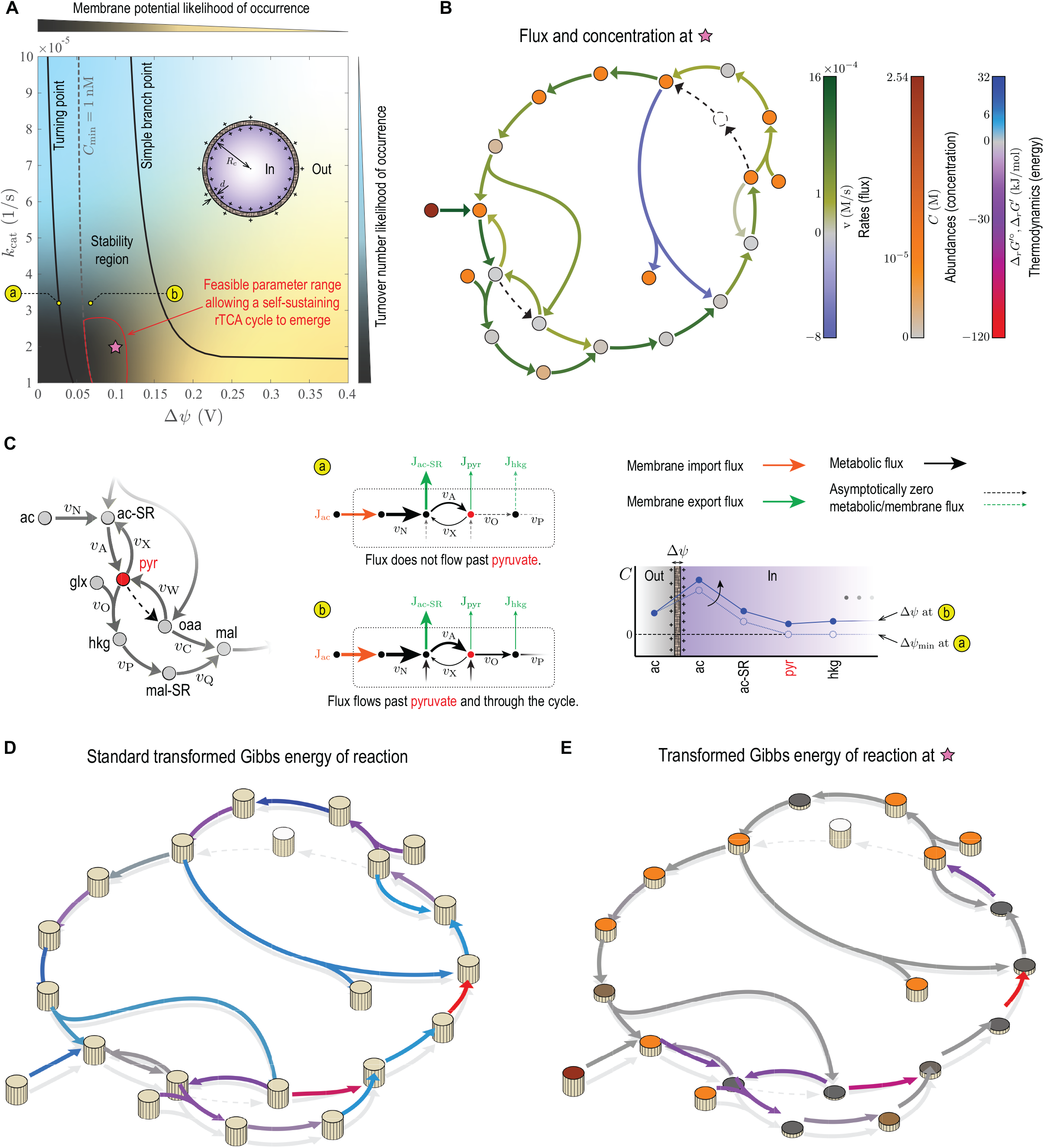
Feasibility of first carbon-fixing cycles at the origin of metabolism. (A) Stability region of the rTCA cycle in Fig. 1C with respect to the turnover number *k*_cat_ and membrane potential Δ*ψ*. The turnover number of all the steps in the network are identical (see Supplementary Information). Steady-state solutions of mass-balance equations are stable in the area confined by solid black lines, where solution branches bifurcate. The dashed line corresponds to a lower bound (1 nM) imposed on the minimum concentration of reactants in the network *C*_min_, splitting the stability region into two subregions. The minimum concentration is less than the lower bound in the left and more than the lower bound in the right sub-region. Generating large membrane potentials (Δ*ψ* 0.15 V) in abiotic cells would have been unlikely due to thermodynamic limitations on charge densities that can be induced on mineral surfaces and stability constraints [18]. Hydrothermal minerals would likely have furnished small turnover numbers (*k*_cat_ ≈ 10^−5^ 1/s) due to their poor catalytic efficiency. The region enclosed by red lines indicates parameter ranges that allow a self-sustaining rTCA cycle to operate. (B) Metabolic fluxes and concentrations at a point inside the stability region that represent likely conditions of hydrothermal vents in the Hadean ocean. (C) Distribution of metabolic fluxes and concentrations near the network entry, where acetate feeds into the rTCA cycle. A minimum membrane potential is required to sustain a stable flux through the entire cycle. Positive membrane potentials generate a large concentration gradient at the cycle entry by elevating the acetate concentration while minimizing the dissipation of downstream intermediates, thus driving a flux through all the thermodynamic bottlenecks. (D) Standard transformed Gibbs energy of reactions. (E) Transformed Gibbs energy of reactions at the same point as fluxes and concentrations in (B) are evaluated. Note that the transformed Gibbs energy determines the spontaneity of reactions when pH is a control parameter besides temperature and pressure [30], as is the case in this work. Model parameters are given in Table S3.

The stability-region boundaries correspond to critical points where multiple steady states exist. At these points, steady states become unstable and solution branches bifurcate. The boundary at low membrane potentials corresponds to turning points. Here, there is a minimum membrane potential at a fixed turnover number below which no steady-state solution associated with a complete-cycle flux exists. Below this minimum limit, the membrane potential is not sufficiently strong to create a large acetate concentration at the network entry to drive a flux through the entire cycle (Fig. 2C). In contrast, the boundary at high membrane potentials corresponds to branch points, where steady-state solutions associated with a complete-cycle flux at a fixed turnover number can be continued along two distinct directions, although these solutions are unstable. Overall, the protocell model exemplifies evolvable systems with coexisting stable and unstable states that are amenable to nongenetic selection [24, 25]. In these systems, physico-chemical properties that yield the most stable states are those that are selected for in long evolutionary processes.

Achievable parameter values under hydrothermal conditions determine part of the stability region that is relevant to the emergence of first metabolic cycles. For example, turnover numbers furnished by hydrothermal minerals would likely have been significantly lower than those of enzymes. Similarly, membrane potentials that were larger in magnitude than those of modern cells would unlikely have been attained due to thermodynamic and stability constraints [18]. Moreover, concentrations should have remained above a minimum level to support metabolic reactions. Accounting for all these restrictions, we qualitatively identified a feasible region (Fig. 2A, area enclosed by red lines) that allows a self-sustaining rTCA cycle to emerge. The resulting parameter ranges are relatively narrow, providing a clue to why it would have been challenging for carbon-fixing cycles to emerge on the early Earth.

The feasibility of carbon-fixing pathways is largely determined by their energetics. The protometabolic network that we examined has several thermodynamic bottlenecks under standard conditions (Fig. 2D, blue arrows). To overcome these energy barriers, a large concentration differential must be generated between the reactants and products of uphill reactions. All concentrations must remain above a minimum level to ensure that a stable flux passes through the entire cycle. To this end, the concentration and import flux of acetate at the entry must be sufficiently large to compensate for the dissipation of the intermediates along the way. The concentration of the intermediates must be distributed around the cycle such that all uphill reactions are eliminated. Any given point inside the stability region corresponds to a complete-cycle flux (Fig. 2B) with no uphill reactions (Fig. 2E). The intermediates in the upper half of the cycle carry more negative charge than those in the lower half. Therefore, by reducing the dissipation rates of the intermediates, positive membrane potentials amplify fluxes and concentrations in the upper half more than in the lower half.

Our protocell model exhibits a growth-like characteristic once it attains a stable steady state, constantly converting substrates (carbon and energy sources) into organic products. Both substrates and products of the protometabolic network are negatively charged. Hence, positive membrane potentials enhance substrate import and alleviate product dissipation. The net effect is an accumulation of more ions in the protocell and a higher osmotic pressure differential than if no metabolic reaction occurred. Overall, the protocell tends to expand as a result of abiotic metabolism. This is a general characteristic of the protometabolic network, irrespective of parameter values (Fig. 3A,B).

**Figure 3:**
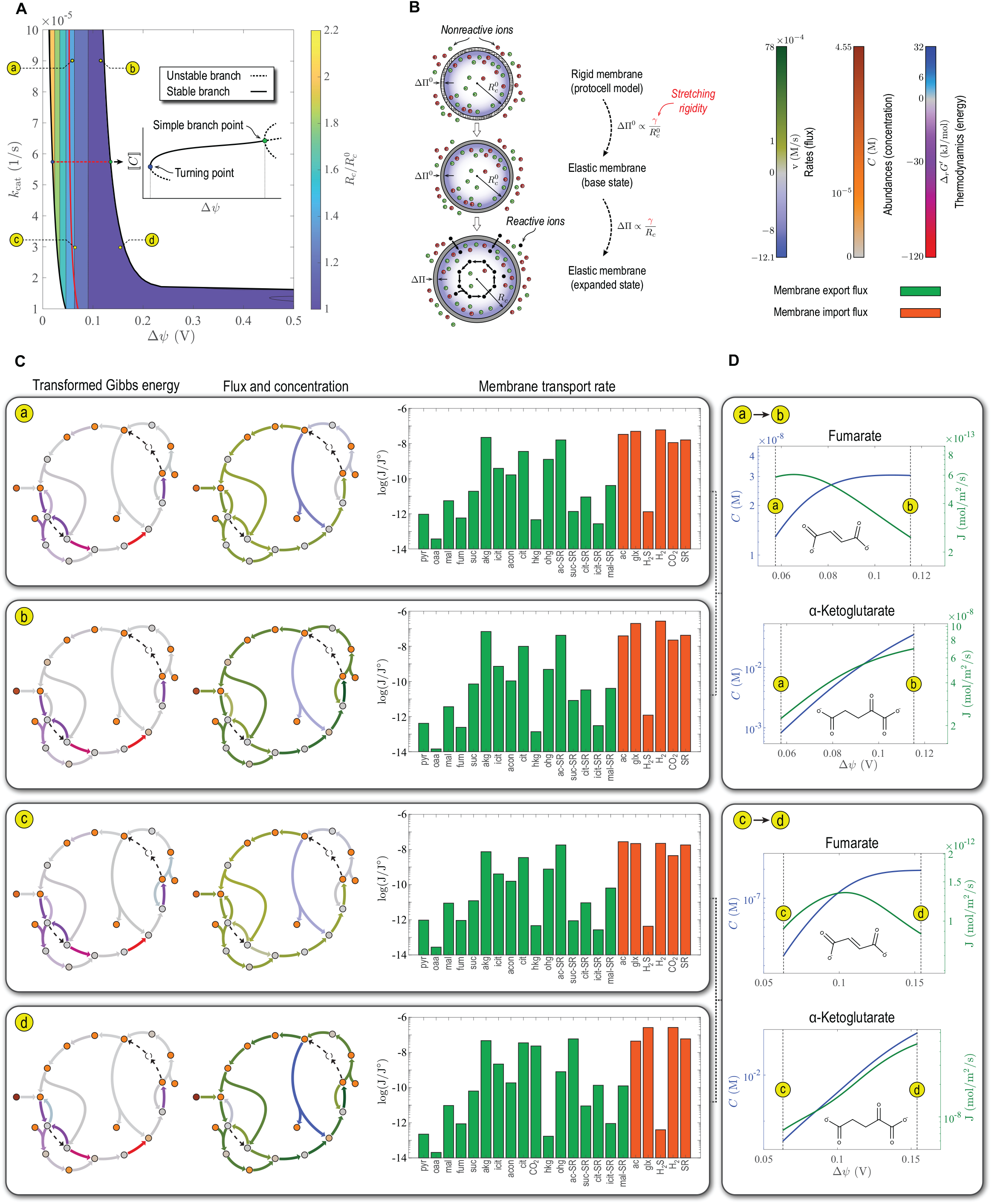
Steady-state characteristics of first carbon-fixing cycles. (A) Volume expansion induced in the protocell by the osmotic pressure resulting from accumulation of the organic products of the rTCA cycle in Fig. 1C. Solid red line corresponds to a minimum-concentration lower bound imposed on steady-state concentrations (dashed line in Fig. 2A). In the inset, ⟦*C*⟧ denotes a solution norm used to represent branching diagrams. (B) Definition of volume expansion shown in (A). The membrane in the protocell model (Fig. 1B) used to characterize the steady states of the rTCA cycle is rigid. However, to estimate the volume expansion resulting from the operation of the protometabolic network, the radius of a protocell with an elastic membrane, containing steady-state concentrations of metabolic products from the protocell model (expanded state), is compared to that of another protocell with an elastic membrane in which only the inorganic ions of the early ocean are present (base state) (see Supplementary Information). The osmotic pressure differential in the first and second case is denoted ΔΠ and ΔΠ^0^ with *R*_*c*_ and 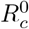 the respective protocell radii. Both metabolic products and inorganic ions contribute to ΔΠ in the first case, while inorganic ions alone generate ΔΠ^0^ in the second case. The stretching rigidity *γ* of the membrane is the same in both cases. (C) Steady-state transformed Gibbs energy of reactions, metabolic fluxes, concentrations, and membrane transport rates at representative points along the boundaries of the stability region. Here, J° = 1 mol/m^2^/s is a reference flux used to nondimensionalize the argument of the logarithm function. (D) Changes in the concentration and membrane transport rate of representative intermediates of the rTCA cycle induced by the membrane potential. Model parameters are given in Table S3.

Both turnover number and membrane potential enhance membrane transport rates and metabolic fluxes. However, the membrane potential has a stronger effect (Fig. 3C), making it the main driving force of metabolism and growth in the protocell model. Moreover, because of nonlinear rate laws and charge-related transport rates, increasing the membrane potential does not affect all the intermediates to the same degree. For example, increasing the membrane potential at large turnover numbers causes the protocell to accumulate and secrete *α*-ketoglutarate at a faster rate than other intermediates (*e*.*g*., fumarate) (Fig. 3D). *α*-ketoglutarate is a precursor for primordial synthesis of glutamate [26]. Glutamate and cysteine in turn can chelate FeS crystals [27, 28], generating a feedback into the reductive carboxylation steps (Fig. S5). Such positive feedbacks could have amplified first carbon-fixing cycles and underlain the evolutionary selection of physicochemical properties that were optimal for amino-acid synthesis.

Here, we demonstrated that iron-sulphide protocells could have harbored first carbonfixing cycles using a first-principle and systems-level approach. These protocells could achieve a sustainable metabolic homeostasis in a narrow parameter range, supporting the hypothesis that, although challenging, cellular metabolism inevitably emerged from geochemical interactions of the early lithosphere and hydrosphere [25]. Stable, self-amplifying carbon fixation was restricted by the membrane potential but unrestricted by the turnover number in our model. These results suggest that, at the origin of metabolism, evolution could have acted on turnover numbers unrestrictedly to improve fitness, maintaining the membrane potential within the tight bounds of physico-chemical constraints. Our first-principle approach could guide future experimental designs to examine the compartmentalization of self-sustaining carbon-fixing networks, opening exciting new directions in the coming years for ‘systems’ studies of life’s metabolic origins.

## Author Contributions

Conceptualization, A.A. and B.O.P.; Methodology, A.A.; Validation A.A.; Formal Analysis A.A.; Investigation, A.A.; Writing – Original Draft, A.A.; Writing – Review & Editing, A.A. and B.O.P.; Funding Acquisition, B.O.P.; Resources, B.O.P.; Supervision, B.O.P.

## Data Availability

All data generated or analyzed during this study are included in this published article and its Supplementary Information files.

## Code Availability

All the codes, their description, and data used for analysis are provided in Supplementary Information files.

## Acknowledgments

This work was funded by the Novo Nordisk Foundation (Grant Number NNF10CC1016517) and the National Institutes of Health (Grant Number GM057089).

## Supporting information

S1 Supplementary Materials: Figures and tables cited in the Materials and Methods section.

## Supplementary Information for

### Model Description

We begin by describing the protocell model that we introduced in the main article to study the emergence of first metabolic cycles. The model we used in the study is similar to the one we previously considered to estimate the membrane potentials that can be generated in primitive cells with iron-sulfide membranes [1]. A primary purpose of this model is to simulate alkaline hydrothermal vent (AHV) conditions on the Early earth. Specifically, the protocell in this model is intended to represent AHV micropores and answer whether they could have harbored early carbon-fixing cycles at the origin of metabolism.

Our model is based on metabolic theories of life’s origins—those that are associated with the metabolism-first hypothesis [2–4]. These theories postulate that life originated from an ancient core metabolic network on the early Earth long before the advent of enzymes or genes. Although what this core network might have been like is unknown, phylogenetic studies have identified some of its characteristics. These studies indicate an anoxic, phosphate-free, and autotrophic network at the core of metabolism [5, 6]. We now have experimental evidence, suggesting that key parts of the core network, including the acetyl-CoA pathway [7, 8], tricarboxylic acid cycle [9–11], glycolysis and pentose phosphate pathway [12, 13], amino-acid and pyrimidine-nucleobase biosynthesis pathways [14, 15], and gluconeogenesis [16] are thermodynamically favored, and they can proceed nonenzymatically. However, whether all or a subset of these nonenzymatic pathways could have spontaneously emerged and formed a self-sustaining metabolic network under plausible prebiotic conditions is yet to be demonstrated.

AHVs in the Hadean ocean would have been a suitable environment for first metabolic pathways to emerge. These environments could have provided steep proton and redox gradients to drive carbon metabolism [17, 18]. Moreover, they would have been rich in CO_2_, reducing gases (*e*.*g*., H_2_ and H_2_S), and transition metal minerals with catalytic and reducing properties [17]. Besides CO_2_, simple C2 carbon sources may also have been present in AHVs. Acetate and glyoxylate are organic C2 carbon sources that could have played an important role at the origin of metabolism. They can be synthesized nonenzymatically from CO_2_ and HCN—prebiotically plausible C1 carbon sources—through the Wood-Ljungdahl pathway [7, 8, 19] and oligomerization reactions [20]. Glyoxylate can also be synthesized from CO—an intermediate of the Wood-Ljungdahl pathway—under hydrothermal conditions using transition metal catalysts without aqueous CN^−^ through the intermediacy of glycolate [21, 22]. These are simple linear pathways that could have operated without enzymes in unconfined environments, such as oceanic crusts (Fig. S1A). However, because of the concentration problem, complex carbon-fixing cycles likely required a confined environment to emerge, stabilize, self-amplify, and evolve. The concentration problem arises as a result of the restrictions associated with nonenzymatic origins of metabolism. Before enzymes, inorganic catalysts (*e*.*g*., transition metals) could have promoted early metabolic reactions. However, because of the poor efficiency of inorganic catalysts, a stable flux through early metabolic pathways could not have been sustained. As a result, metabolic fluxes could have hardly counterbalanced diffusive losses in unconfined environments to concentrate the intermediates of these pathways beyond a minimum physical limit [1, 23, 24]. Although the concentration problem affects both linear and cyclic pathways, it would have limited the feasibility of cyclic pathways more [25]. Hence, it is plausible to assume that first carbon-fixing cycles emerged in confined regions of AHVs in the primitive ocean (Fig. S1A).

Given the importance of confinement in the emergence and evolution of complex metabolic networks, transition to cellular metabolism from very early stages of chemical self-organization might have been inevitable. In the context of the metabolism-first hypothesis, abiotic cells that contained first metabolic cycles must have been made of purely inorganic materials, and all metabolic reactions in these cells must have been promoted by inorganic catalysts without relying on enzymes. In the absence of enzymes, the operation of abiotic cells would have been restricted in several ways. For example, the import of energy or carbon sources and export of products should have occurred unselectively through passive diffusion. Carbon fixation should have been driven by primitive energy-coupling means without electron bifurcation or proton pumps. Moreover, reactant concentrations and metabolic fluxes would have been extremely low due to the poor efficiency of inorganic catalysts, limiting the feasibility of primitive pathways, especially those with multiple thermodynamic bottlenecks.

**Figure S1:**
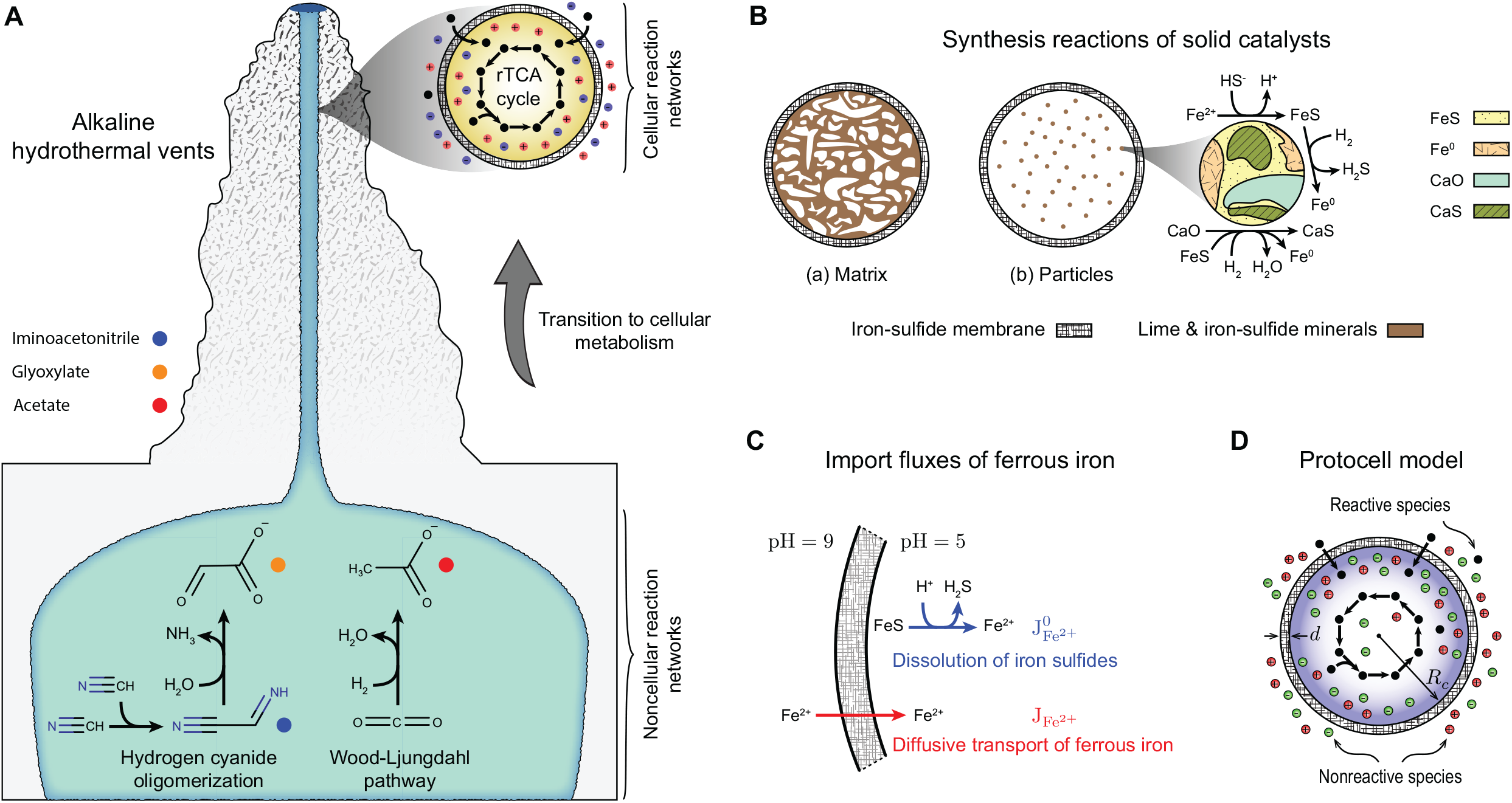
Cellular origins of metabolism. (A) AHVs could have harbored earliest cellular structures capable of sustaining carbon-fixing cycles on the early Earth. C2 organic compounds, such as acetate and glyoxylate, could have been synthesized in deep seabeds from inorganic C1 carbon sources, such as CO_2_ and HCN. Acetate and glyoxylate could have been produced nonenzymatically through the Wood-Ljungdahl pathway [7, 8, 19] and HCN-oligomerization reactions [20] under prebiotic conditions. Because these pathways are linear and simple, they could have sponteneously arisen in oceanic crusts in unconfined environments. However, autocatalytic carbon-fixing pathways would likely have required a controlled environment to operate and self-amplify, necessitating a transition to cellular metabolism. Thin-walled vent micropores made of iron sulfides could have furnished a suitable environment for first carbon-fixing cycles to emerge, providing a barrier to facilitate primitive forms of energy coupling and contain the organic products of the foregoing cycles. (B) The protometabolic network in Fig. 1C is driven by inorganic catalysts and reducing agents, such as FeS and Fe^0^. Under hydrothermal conditions, mixtures of FeS and Fe^0^ can be synthesized through the interaction of Fe^2+^ and HS^−^ [26]. As a result of these interactions in the protocell model, assemblages of FeS-Fe^0^ precipitate on lime minerals, which would have been present in hydrothermal-vent environments. The resulting mineral mixtures are distributed inside the protocell either as a continuous matrix (a) or dispersed particles (b). (C) Ferrous iron that participates in metabolic reactions and mineral synthesis enters the protocell from outside by passive diffusion. The dissolution of the iron-sulfide membrane on the acidic side also yields ferrous iron, which contributes to its import flux. The dissolution rate of iron-sulfide membrane and the membrane transport rate of ferrous iron are denoted 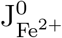 and 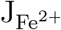, respectively. (D) Protocell model, simulating the operation of first carbon-fixing cycles in AHVs micropores.

AHV micropores could have served as the earliest cellular structures in which first carbon-fixing cycles emerged. However, whether a self-sustaining carbon-fixing cycle can operate on purely inorganic precursors (*i*.*e*., energy sources, reducing compounds, and catalysts) alone in such inorganic cells is unclear. To answer this question, we developed a protocell model of AHV micropores. Our goal is to demonstrated the feasibility of such protocells despite the highly restrictive physico-chemical constraints of the early Earth.

Since fully autotrophic pathways would have been challenging to operate without enzymes, we study the feasibility of partially autotrophic carbon fixation by including C2 carbon sources, such as acetate and glyoxylate, in the food set. As previously stated, these could have been synthesized from inorganic C1 carbon sources in oceanic crusts of the primitive Earth under prebiotic conditions. In particular, because the partial pressure of CO_2_ in hydrothermal systems of the early Earth would have been significantly higher than those in modern oceans [17, 18, 27], acetate could have been synthesized in high concentrations (*C* ≈ 1 M) through the Wood-Ljungdahl pathway. Acetate and glyoxylate, which were present in AHV environments, then would have been transported into primitive cells along with CO_2_ by passive diffusion (see Fig. 1B).

We consider an idealized protocell model of AHV micropores. The protocell is a sphere of radius *R*_*c*_ with a semi-permeable membrane of thickness *d* (Fig. S1D). The membrane is primarily made of iron sulfides possibly mixed with other minerals. Under hydrothermal conditions, similar cellular structures made of iron monosulphide can form when ferrous iron is mixed with alkaline fluids that are rich in HS^−^ [28]. We assume that a simple hypothetical thiol HS–R (*e*.*g*., ethane thiol [17]) existed in primitive AHVs that drove the thioesterification steps of first metabolic networks. The interior of the protocell is filled with acidic ocean water (pH = 5), while its outer membrane is exposed to alkaline hydrothermal fluids (pH = 9) (see Fig. 1A,B). Because of this pH gradient, a positive membrane potential Δ*ψ* develops across the membrane [1]. Hereafter, we refer to the acidic interior of the protocell as the intracellular environment (denoted ‘In’ in Fig. 2A) and alkaline hydrothermal fluids outside the protocell as the extracellular environment (denoted ‘Out’ in Fig. 2A). Carbon sources, energy sources, and all the organic intermediates of first metabolic cycles in our model are negatively charged. Thus, positive membrane potentials facilitate the import of carbon and energy sources, while alleviating the dissipation of the organic intermediates. The protocell imports carbons sources (acetate, glyoxylate, and CO_2_), reducing gases (H_2_ and H_2_S), and thiols (HS–R) from outside. It has no specialized ion channels or transporters, so the only mechanism of membrane transport is unselective passive diffusion subject to an electric field.

Except carbon-fixing steps, all metabolic reactions are catalyzed by transition metals in aqueous (Fe^2+^ and Cr^3+^) or solid (Fe^0^ and FeS) phase. They are dispersed inside the protocell in an acidic environment. Beside iron sulfides, lime minerals would likely have been available in AHVs of the primitive ocean. Accordingly, we assume that the protocell is filled with lime minerals, which either form a continuous matrix (Fig. S1B(a)) or dispersed particles (Fig. S1B(a)). Assemblages of FeS-Fe^0^ precipitate on these lime minerals upon interaction of Fe^2+^ and HS^−^ ions (Fig. S1B). Interestingly, these assemblages exhibit superior catalytic efficiencies to pure FeS in promoting prebiotically important reduction, reductive carboxylation, and reductive amination reactions [26]. In our protocell model, several steps of the protometabolic network that we examined in this work are catalyzed by these mineral mixtures (see “Kinetic Constraints”). On the other hand, the reductive carboxylation steps of the network are driven by protoferredoxins. We assume that the reduced and oxidized forms of these protoferredoxins, respectively denoted FD_rd_ and FD_ox_, constitute a fraction of the iron-sulfide membrane. The energy released by the redox conversion FD_rd_ → FD_ox_ drives the reductive carboxylation steps on the inner surface of the membrane in an acidic environment. On the outer surface of the membrane, H_2_ reduces FD_ox_ in an alkaline environment to regenerate FD_rd_. Note that reductive carboxylation and FD_rd_-regeneration reactions are more thermodynamically favorable in acidic and alkaline environments, respectively.

**Figure S2:**
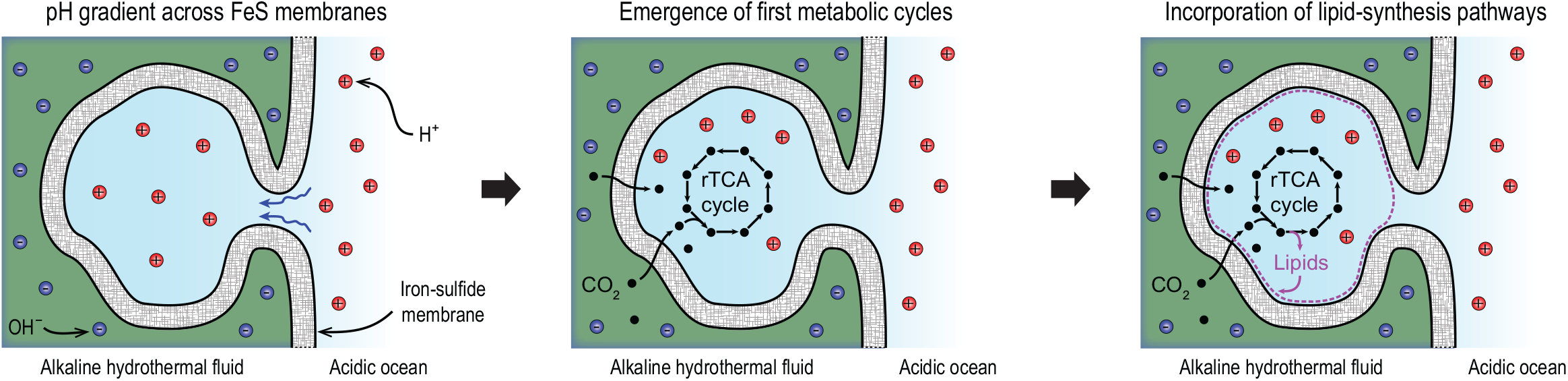
Maintaining pH gradient across the membranes of earliest cellular structures. As suggested in previous studies [29], thin-walled AHV cavities made of iron sulfides could have served as inorganic cells, harboring the first metabolic networks at the origin of metabolism. The inorganic membranes of these cellular structures could have separated alkaline hydrothermal fluids from acidic ocean waters. Early metabolic networks could have incorporated lipid-synthesis pathways at later stages once their precursors had been produced in sufficiently large concentrations. Single-chain amphiphiles produced by these pathways could have self-assembled into lipid membranes [30], replacing iron-sulfide membranes.

Acidic environments can dissolve iron sulfides. In our protocell model, the rate at which Fe^2+^ is released into the protocell as a result of the dissolution of the iron-sulfide membrane 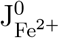 constitute part of the import flux of Fe^2+^. The other part 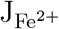 is due to its diffusive transport from the extracellular environment (Fig. S1C). The ferrous iron entering the protocell through these import fluxes interact with HS^−^ ions to form the foregoing FeS-Fe^0^ assemblages. Other metal ions involved in metabolic reactions (*e*.*g*., Cr^3+^) are also imported from the extracellular environment similarly to Fe^2+^.

Note that the idealized protocell model shown in Fig. S1D represents both closed and open AHV micropores. It is intended to capture important physico-chemical characteristics of hydrothermal systems that played a role in the emergence of first metabolic networks on the early Earth. In the main article, we provided a schematic representation of closed micropores (Fig. 1A, inset) as abiotic cells that could have harbored first carbon-fixing cycles at the origin of metabolism. However, open cavities in AHVs, such as that shown in Fig. S2, have also been considered in previous studies on the origins of metabolism [29]. These cavities could have maintained a pH gradient across their inorganic membranes to drive carbon fixation. Once stabilized, early carbon-fixing cycles could have produced the precursors of fatty acids, incorporating lipid-synthesis pathways [1, 30]. Lipid membranes could have spontaneously emerged in such cavities from the self-assembly of single-chain amphiphiles produced by these pathways [30, 31] and replaced iron-sulfide membranes at later stages (see Fig. S2).

### First Carbon-Fixing Cycles at the Origin of Metabolism

According to metabolic theories of life’s origins, life emerged as a consequence of the natural tendency of a self-organized geochemical reaction network to dissipate free energy [17, 32]. This network would have been thermodynamically favored and kinetically assisted by mineral catalysts. Thus, it could have occurred spontaneously and operated without enzymes. Such a protometabolic network would not have been fundamentally different than biochemical reactions at the core of biological metabolism in substrates, topology, catalysts, or energy-coupling mechanisms [33]. Given the highly interconnected structure of reaction networks leading up to extant biology and their nonlinear dynamics, a major change in these characteristics would likely have resulted in a catastrophic event and a sudden transition to a chaotic and nonfunctional system. There has been no evidence of such drastic transitions in the evolutionary history of metabolism at least as far back as 3.5 billion years ago, and they were unlikely to have had happened before [32]. Therefore, it is reasonable to search for metabolic origins that are congruent with modern metabolism.

All essential metabolic processes of life hinge on carbon fixation in some form. It is an important biosynthesis route for building complex molecules from simpler ones, and it would likely have been the primary function of the earliest metabolic pathways. Thus, a long-standing question in prebiotic chemistry has been whether there were primitive reaction networks at the origin of metabolism that could have fixed CO_2_—one of the most abundant inorganic carbon sources on the early Earth [34]. Using life as a guide to its origins, one may search for clues to the chemistry underlying these reaction networks in the structure of CO_2_-fixing pathways in modern autotrophs. From the six know CO_2_-fixing pathways, the acetyl-CoA pathway and the reductive tricarboxylic acid (rTCA) cycle are ancient pathways that could provide insight into what the first CO_2_-fixing networks were like [3, 25, 34]. The acetyl-CoA pathway is present in both bacteria and archaea [29], while the full rTCA cycle is only found in bacteria [32]. They both could have played a significant role in the emergence of metabolic pathways. However, whether the complete rTCA cycle or its other forms could represent the underlying chemistry of the first carbon-fixing pathways has not been established. Nevertheless, a hybrid network comprising the acetyl-CoA pathway and the horseshoe rTCA cycle (*i*.*e*., pyruvate to *α*-ketoglutarate) [32], a pyrite-pulled rTCA-cycle analogue [34], and a pyrophosphate driven biology-like rTCA cycle [35, 36] have been proposed as plausible models of prebiotic carbon fixation.

Although the antiquity of the complete rTCA cycle is not fully supported by the phylogeny of modern bacteria and archaea, it is still plausible to assume that some variant of it was the first CO_2_-fixing pathway, or at least played a significant role in one, for several reason [17, 32, 34, 35]: (i) It is a truly autocatalytic cycle, so it can self-amplify and undergo evolutionary selection provided its carbon and energy sources are readily available in the environment. (ii) It is anoxic and fully autotrophic. Once it has been started and stabilized, it fixes four CO_2_ molecules with every turn without needing other carbon sources. (iii) Except cofactors, all its intermediates are phosphate free. (iv) It contains the five universal intermediates from which all the building blocks of life (*i*.*e*., amino acids, fatty acids, and nucleotides) are biologically synthesized. (v) So far, transition metals have been shown to promote seven out of thirteen steps of the cycle [9, 37]. (vi) Several enzymes that catalyze the reactions of the cycle have iron-sulfur clusters or transition-metal binding sites (see Fig. S3).

**Figure S3:**
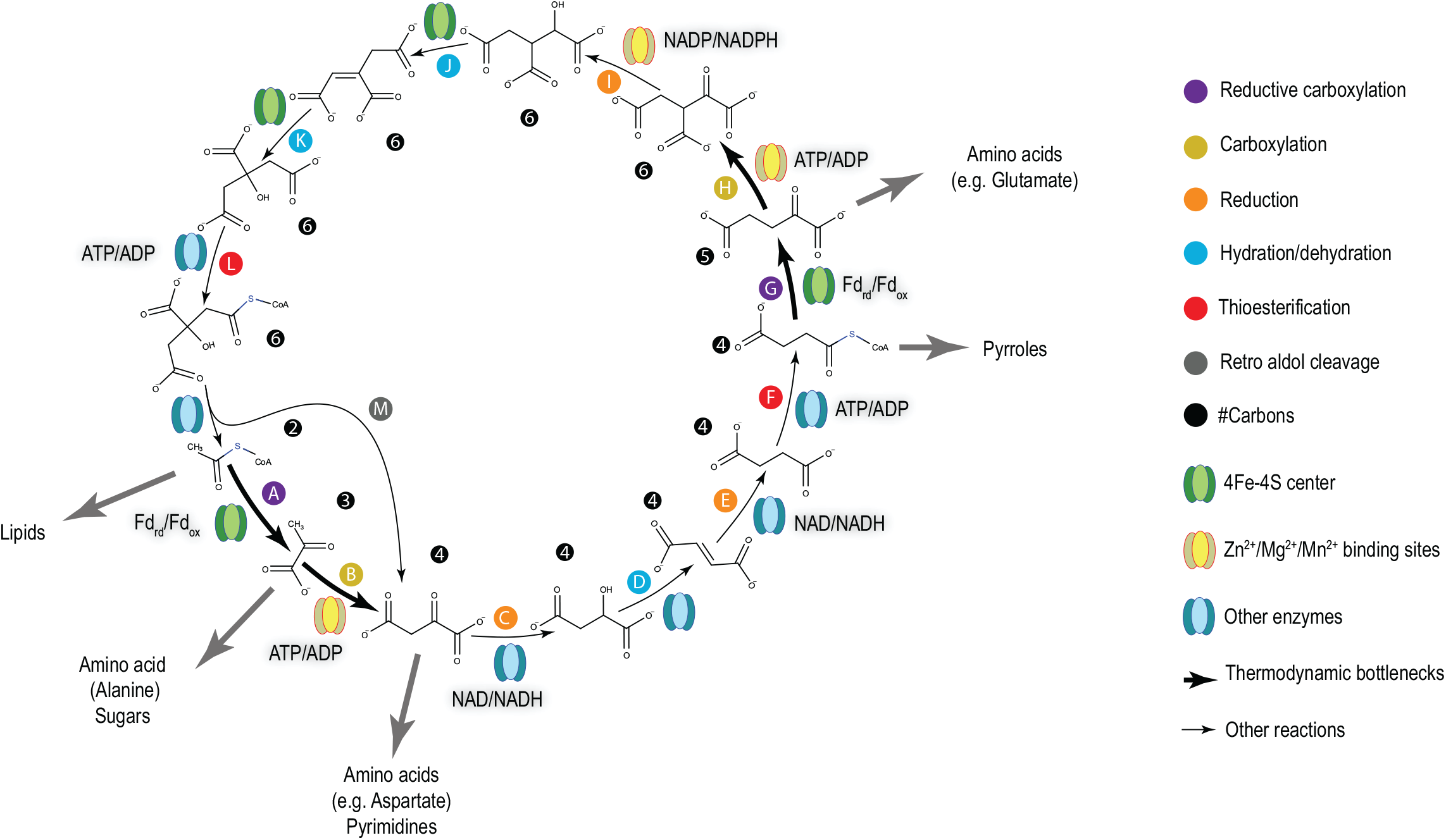
The enzymatic rTCA cycle. This ancient pathway contains the five universal intermediates from which all the building blocks of life, including lipids, amino acids, sugars, pyrimidines, and pyrroles are synthesized. The cofactors driving reduction or energy-demanding steps are highlighted in gray.

The enzymatic rTCA cycle may be an ancient biochemical reaction network. It could have been present in the Last Universal Common Ancestor, but part of it might have been lost in archaea. Nevertheless, the biological rTCA cycle is not an accurate representation of prebiotic CO_2_-fixing cycles: First, several reactions in the enzymatic rTCA cycle are driven by phosphate-containing cofactors, such as ATP and NADH. However, some of these steps can also be promoted nonenzymatically by transition metals [9, 10]. These metal-catalyzed reactions might have preceded their biological counterparts before the emergence of enzymes. Second, the thioesterification steps of the rTCA cycle are facilitated by ATP and coenzyme A. Both contain multiple phosphate groups and are products of a long evolutionary process [17], so they could not have played any role at the origin of metabolism. However, thioesterification can also be accomplished by simple, inorganic thiols with acids or metal ions as catalysts [38, 39]. Third, the carbon-fixation steps of the rTCA cycle are highly endergonic. These thermodynamic bottlenecks must be overcome to achieve a self-sustaining complete-cycle flux state. Modern organisms (*e*.*g*., thermophiles) can sustain a complete rTCA cycle thanks to their efficient and evolutionary optimized enzymes [40]. However, in the absence of enzymes, the alternative inorganic catalysts would not have been efficient enough to diminish the kinetic barriers associated with the thermodynamic bottlenecks. Consequently, maintaining a fully cyclic flux in early metabolic pathways without enzymes would have been extremely challenging. Hence, energy-demanding reactions should have been coupled to inorganic energy sources by prebiotic mechanisms.

**Figure S4:**
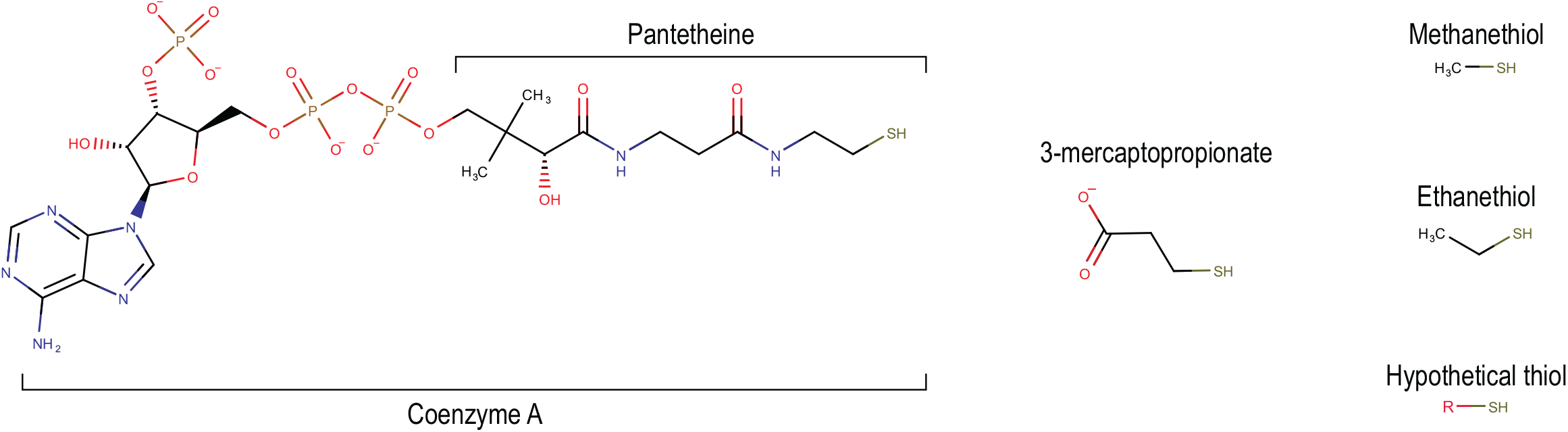
Thiols participating in enzymatic and nonenzymatic thioesterification reactions. The thioesterification steps of the protometabolic network in this work are driven by a simple, hypothetical thiol that could have been present in the hydrothermal vents of the Hadean ocean with R a hypothetical substituent.

Addressing the foregoing three problems, we derived a protometabolic network from the rTCA cycle (Fig. 1C) that is compatible with the AHV conditions and could have operated prebiotically. Firstly, we replaced all enzymatic reactions that can be promoted by transition metals with their nonenzymatic counterpart. Specifically, in our protometabolic network, hydration and dehydration steps are catalyzed by Fe^2+^ and Cr^3+^, while reduction steps are promoted by Fe^0^. As discussed in section “Model Description”, Fe^2+^ and Cr^3+^ are imported into the protocell from the extracellular environment, and Fe^0^ is synthesized through the interaction of Fe^2+^ with HS^−^.

Furthermore, coenzyme A is replaced by a hypothetical thiol HS–R in the thioesterification steps of the protometabolic network. Such simple, inorganic thiols could have been synthesized in the sulphide-rich AHVs of the primitive ocean. Previous studies on core reaction networks preceding phosphate-based metabolism considered simpler phosphate-free variants of coenzyme A, such as pantetheine, 3-mercaptopropionate, methanethiol, and ethanethiol as prebiotic thiols involved in thioester chemistry (see Fig. S4) [5, 17]. In this work, the chemical structure of the substituent R is unspecified. Accordingly, to estimate the transformed Gibbs energy of thioesterification reactions, the transformed Gibbs energy of formation of this hypothetical thiol and its corresponding thioesters are provided as given parameters of the model (see “Abiotic Thioesterification”). However, to reduce the number of model parameters, its diffusivity coefficient is approximated based on the van der Waals radius of ethanethiol (see “Mass Balance Constraints and Maxwell’s first law”).

Finally, we considered primitive energy-coupling mechanisms to overcome the energy barriers associated with carbon fixation. In the enzymatic rTCA cycle, there are four carbon-carbon bond-forming reactions: two reductive carboxylation steps driven by ferredoxin and two neutral carboxylation steps coupled to ATP hydrolysis (see Fig. S3). Among these, only the reductive carboxylation of acetate to pyruvate has been experimentally shown to occur without enzymes using iron-containing minerals (*e*.*g*., greigite, magnetite, and awaruite) as catalysts and hydrogen as the reducing agent [8]. Therefore, it is plausible to assume that, in early metabolic cycles, both reductive carboxylation steps were promoted by primitive redox couples, possibly comprising iron, nickle, and sulfur with a similar crystal structure to Fe-S clusters in modern ferredoxin [17]. However, the neutral carboxylation steps have not been reproduced nonenzymatically in laboratory experiments so far, and the were unlikely to have occurred under prebiotic conditions without evolutionarily optimized enzymes or cofactors, such as ATP.

In the remainder of this section, we elaborate further on the four carbon-fixation steps discussed above. First, consider the reductive carboxylation steps. In our protocell model, the two reductive carboxylation steps are driven by protoferredoxins. In extant biology, these steps are accomplished by ferredoxins—enzymes containing Fe_4_S_4_ thiocubane centers that are ligated to proteins by four cysteine residues. Strong metal-ligand bonds stabilize the crystal structure of Fe-S clusters in these enzymes [41]. The crystal structure in turn determines the reduction potential and catalytic capacity of Fe-S clusters, although other factors might also play a role [42]. Moreover, the ligands mitigate oxidative degradation of Fe-S clusters in the presence of oxidizing agents, such as Brønsted acids or hydrogen peroxide [43]. What forms of protoferredoxins could have been synthesized under prebiotic conditions? In the absence of proteins, could naturally occurring hydrothermal redox couples have instead driven the reductive carboxylation steps of early metabolic networks?

Although hydrothermal redox couples such as mackinawite/greigite and pyrrhotite/pyrite can promote carbon-fixation reactions similarly to those in acetogens [8] and methanogens [44], whether they could have played a significant role at early evolutionary stages of metabolism is in question. Pyrite is a highly stable form of iron sulfides with a complex crystal structure, making it hard to reduce. Once it is produced, it can no longer participate in abiotic metabolic reactions, nor can it be regenerated by reducing agents, such as H_2_ or H_2_S [17]. Regenerability is an important feature of extant redox couples (*e*.*g*., NAD/NADH and FAD/FADH), underlying the autonomy and robustness of modern organisms. In contrast, mackinawite and greigite are electroconductive with readily switchable iron valence; thus, in principle, mackinawite can be regenerated by reducing greigite [17]. Interestingly, the crystal structure of greigite resembles that of iron-sulfur centers in modern ferredoxins, so it is expected to exhibit similar catalytic characteristics [17]. However, both amorphous FeS and greigite are prone to oxidative degradation. That is, they are both oxidized to pyrite if exposed to H_2_S and H^+^ [45]. This last point is particularly problematic in the context of autotrophic origins of metabolism: On the one hand, acidic environments thermodynamically favor reductive carboxylation. On the other hand, they degrade the crystal structure of mackinawite and greigite. Therefore, preserving the structural integrity and catalytic capacity of hydrothermal minerals through oxidation and reduction cycles under conditions that allowed abiotic metabolism to spontaneously emerge would have been challenging on the early Earth, necessitating the formation of stabilizing metal-ligand bonds.

Comparative studies of amino-acid sequences of ferredoxins across diverse organismal groups suggest that these archaic proteins have evolved through a sequence-doubling process, whereby their stability and catalytic activity were improved in a stepwise manner [46]. This process started from a four amino-acid peptide and continued by repeating longer and longer sequences of amino acids. Extrapolating this process to the most ancient ferredoxins, it follows that the earliest redox-active iron-sulfur assemblages at the origin of metabolism, which we refer to as protoferredoxins, were stabilized by primitive, short ligands that would have been present on the early Earth. The organic intermediates of the rTCA cycle (*e*.*g*., pyruvate) [45] and prebiotic thiolates [17] are examples of such ligands that could have stabilized primitive Fe-S clusters. Ancient peptides could also have played the same role. As has been experimentally demonstrated, in the presence of iron and sulfur ions, micromolar concentrations of ancient peptides (*e*.*g*., cysteine and glutathione) can chelate redox-active Fe-S clusters [43, 47]. Their constituent amino acids in turn can be synthesized nonenzymatically through metal-catalyzed reductive amination or transamination of *α*-keto acids of the rTCA and glyoxylate cycle under prebiotic conditions [10, 14, 48, 49]. The formation of these amino acids hinge on abiotic carbon-fixing cycles that can produce their precursors in sufficiently large concentrations. To avoid the circular interdependence between amino-acid synthesis and carbon fixation, reductive amination and transamination reactions are assumed not to be activated in the earliest carbon-fixing cycles represented by our protometabolic network, although they could have become activated at later stages. Accordingly, protoferredoxins in our model are synthesized from either the intermediates of the rTCA cycle or primitive thiolates. These protoferredoxins are mixed with other hydrothermal redox couples (*e*.*g*., mackinawite/greigite), and they constitute a fraction of the iron-sulfide membrane of the protocell (see Fig. 1B). As previously stated, these hydrothermal minerals could have promoted the reductive carboxylation steps of early metabolic networks. However, in the protometabolic network, these steps are assumed to be driven only by protoferredoxins for simplicity.

**Figure S5:**
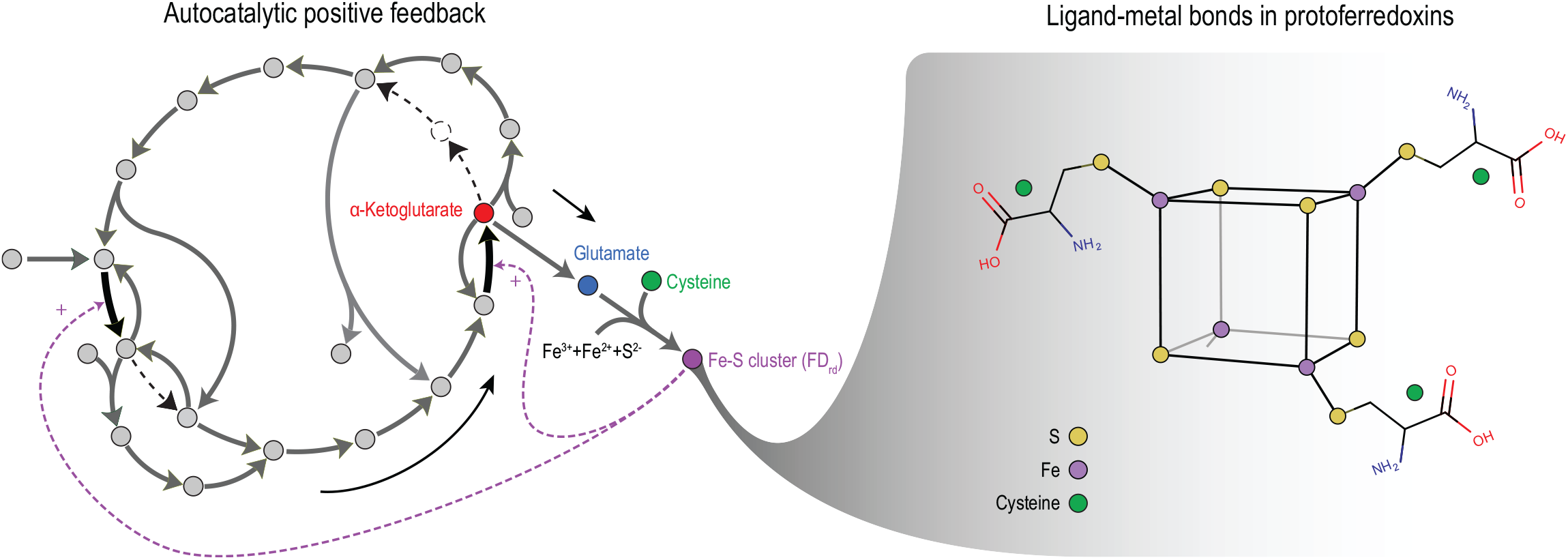
Autocatalytic positive feedback underlying the evolutionary selection of protoferredoxins and amino-acid synthesis pathways in first metabolic cycles. Once their synthesis pathways had been activated, ancient amino acids, such as cysteine, could interact with the products of reductive carboxylation reactions, to chelate Fe-S clusters in the presence of iron and sulfur ions, synthesizing protoferredoxins. The crystal structure of these protoferredoxins is more stable and their catalytic efficiency is higher than those of naturally occurring hydrothermal minerals, such as mackinawite and greigite. Protoferredoxins in turn can accelerate reductive carboxylation reactions more efficiently than alternative mineral catalysts. Thus, the formation of protoferredoxins could have generated positive feedbacks by amplifying the reductive carboxylation steps of first metabolic cycles—a nongenetic mechanism underlying their evolutionary selection.

Autocatalysis could have played an important role in the evolution of first metabolic cycles, such as our protometabolic network, before the emergence of genetic codes [50]. Once these cycles had stabilized, they could produce their intermediates like *α*-ketoacids in large concentrations due to their self-amplifying characteristics. Sufficiently large concentrations of *α*-ketoacids then could have activated amino-acid synthesis pathways in the presence of nitrogen sources. As stated above, ancient amino acids, such as cysteine and glutamate, could have facilitated the formation of more stable and efficient protoferredoxins than those chelated by the intermediates of the rTCA cycle or primitive thiolates. These protoferredoxins in turn could have generated positive feedback, amplifying reductive carboxylation reactions (Fig. S5). Such positive feedback likely underlay the evolutionary selection of amino-acid synthesis pathways and iron-sulfide crystal structures that were optimal for the operation of first carbon-fixing cycles before genes.

Next, consider the neutral carboxylation steps. Without neutral carboxylation, some intermediates of the enzymatic rTCA cycle cannot be accessed. In the absence of energetic cofactors of extant biology, abiotic carboxylation in primitive pathways would have been highly endergonic and challenging to run nonenzymatically. Thus, a natural question is whether nonenzymatic carbon-fixing pathways similar to the rTCA cycle could have operated prebiotically using inorganic energy sources and reducing agents—pathways that were fully cyclic and contained most or all the intermediates of the enzymatic rTCA cycle. Examples of such pathways were examined in the glyoxylate scenario [51]. The pathways considered in this scenario rely on the chemistry of HCN. Glyoxylate is the central intermediate of these pathways, realizing aldol condensations that connect simple C1 carbon sources to more complex intermediates of the rTCA cycle. A recent theoretical analysis showed that a phosphate-free core metabolic network enriched with major carbon-fixing cycles can emerge from simple carbon sources using network expansion algorithms [5]. Interestingly, glyoxylate and pyruvate were among the largest branching points of this core network, suggesting their prominent role in the earliest carbon-fixing pathways. Subsequent experiments confirmed that an abiotic reaction network, comprising a large portion of the TCA and glyoxylate cycle, can be generated using glyoxylate and pyruvate [10]. This network was promoted by Fe^2+^ and produced nine of the thirteen intermediates of the rTCA cycle.

The foregoing studies highlight the importance of glyoxylate for prebiotic production of the intermediates of the rTCA cycle—primordial hubs from which the earliest building blocks of life were likely synthesized. Thus, we searched for nonenzymatic and fully cyclic carbon-fixing pathways among the reactions of the core metabolic network that have been previously demonstrated to proceed spontaneously only using transition metals as catalysts [9, 10]. We considered pathways that could bypass the neutral carboxylation steps to connect simple carbon sources, such as CO_2_ or acetate, to the intermediates of the rTCA cycle if glyoxylate was included in the food set. We found that both neutral carboxylation steps of the enzymatic rTCA cycle can be bypassed through aldol reactions involving glyoxylate followed by oxidative decarboxylation and thioester hydrolysis steps. Specifically, the carboxylation of pyruvate to oxaloacetate (Fig. S3, step B) and of *α*-ketoglutarate to oxalosuccinate (Fig. S3, step H) can be bypassed through aldol reaction of glyoxylate with pyruvate (Fig. 1C, step O) and *α*-ketoglutarate (Fig. 1C, step R), respectively. These off-cycle reactions yield two intermediates that are not produced by the enzymatic rTCA cycle, namely hydroxyketoglutarate and oxalohydroxyglutarate [10]. These intermediates in turn can undergo an oxidative decarboxylation and a thioester hydrolysis to produce malate and isocitrate, respectively. Note that Fe^2+^ can also promote the direct synthesis of malate and isocitrate from hydroxyketoglutarate and oxalohydroxyglutarate without thioester hydrolysis steps if thiols are not present in the reaction environment [10]. However, it is likely that the foregoing bypass pathways occurred predominantly through thiol-mediated oxidative decarboxylation because the alkaline hydrothermal environments of the primitive ocean would have been rich in sulfide and thiols. Accordingly, we neglect the direct decarboxylation reactions in our protometabolic network for simplicity (see Fig. 1C).

### Abiotic Thioesterification

Thioesters play a prominent role in biochemistry, especially in the bioenergetics of metabolic networks [23]. Being coupled to ATP synthesis and energy-demanding condensation reactions [32], they are involved in both catabolic and anabolic processes. In the context of carbon assimilation, thioesters are high-energy and activated forms of carboxylic acids. Specifically, in the rTCA cycle, their synthesis precedes the carboxylation steps to facilitate energy-demanding C-C bond formations. In biological systems, thioesterification is coupled to ATP hydrolysis because it is an endergonic process. However, thioesterification can also be accomplished inorganically using Brønsted acids [38] or metal ions [39] as catalysts. As previously stated, thioesterification steps are driven by a hypothetical thiol HS–R in our protocell model. All the thioesterification steps of our protometabolic network can be represented by the following general reaction

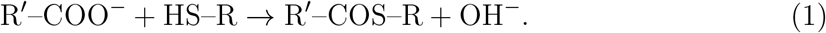

Because the molecular structure of the hypothetical substituent R in this equation is unspecified, the Gibbs energy of formation of HS–R and R^*/*^–COS–R are unknown. To estimate the Gibbs energy of thioesterification reactions in our model, the difference between the reference Gibbs energy of formation of the minimum charge state of each carboxylic acid and its respective thioester, defined by

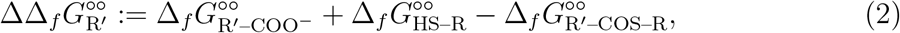

are given as model parameters (Table S1). Here, the superscript ° ° denotes the reference Gibbs energy of formation of the minimum charge state of the species indicated by the subscript. Since these parameters only provide the Gibbs energy of formation of the unspecified thioesters R^*/*^–COS–R relative to that of HS–R, we use the reference Gibbs energy of formation of the minimum charge state of ethanethiol as a basis for the Gibbs energy of formation of HS–R.

In our model, thioesterification reactions are catalyzed by both acids and Fe^2+^. Similarly to biochemical reactions, we assume that the nonenzymatic reactions in our protometabolic network occur at a fixed pH. When pH is a control parameter besides temperature and pressure, the spontaneity of chemical reactions is determined by the transformed Gibbs energy of reaction [52, 53]. This Gibbs energy accounts for acid dissociation constants and depends on the concentration of H^+^ (see “Thermodynamic Constraints”). In this formulation, the catalytic effect of Brønsted acids is captured by the change in the transformed Gibbs energy of reaction because of variations in the acid concentration. According to this formulation, thioesterification reactions have a larger negative transformed Gibbs energy of reaction and are more thermodynamically favorable in more acidic environments [53] because they consume H^+^. As stated before, in our models, thioesterification reactions occur inside the protocell in an acidic environment, where they are both thermodynamically and kinetically favored. Accordingly, protocells with an acidic interior would have provided a desirable environment for first carbon-fixing cycles to emerge by alleviating their thermodynamic bottlenecks.

**Table S1:** Parameters used to estimate the Gibbs energy of formation of hypothetical thioesters in the protometabolic network.

| Carboxylic acid | Identifier | R' | $\Delta\Delta_f G_{R'}^{\circ\circ}$ (kJ/mol) |
| --- | --- | --- | --- |
| Acetic acid | ac | CH3 | -161.5 |
| Succinic acid | suc | COOCH2CH2 | -152.0 |
| Citric acid | cit | COOCH2COHCOOCH2 | -161.5 |
| Isocitric acid | icit | COOCH2CHCOOCHOH | -161.5 |
| Malic acid | mal | COOCHOHCH2 | -142.5 |

Once the transformed reference Gibbs energy of formation of all the species in Eq. (1) and their acid dissociation constants have been determined, the transformed Gibbs energy of the thioesterification steps at a given pH can be ascertained. Note that, besides the reference Gibbs energy of formation of the minimum charge state, we used the acid dissociation constants of ethanethiol to estimate those of the hypothetical thiol in our model.

### Mass Balance Constraints and Maxwell’s first law

Conservation of mass is a fundamental constraint that we impose in our protocell model. We solve the mass-balance equations to determine how the concentrations of different compounds in the protometabolic network vary with time. Before proving the detailed formulation of these equations, we introduce a few concepts that are important in the forthcoming discussion. One is the idea of combining the protonation states of a molecule into an aggregate compound, as defined by Alberty [52]. Most molecules that participate in protometabolism behave like weak acids in water, releasing hydrogen ions in multiple steps

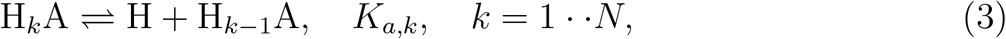

where *N* is the number of hydrogen-dissociation steps, and A represents the minimum charge state of a molecule in the system with *K*_*a,k*_ the respective hydrogen-dissociation constants. These protonation or charge states are referred to as species [52, 53]. Their distribution can vary depending on the pH and ionic strength of the solution. In general, all these charge states are utilized by metabolic networks, so hydrogen-dissociation reactions must be incorporated into metabolic reactions to account for the effect of pH.

Hydrogen-dissociation reactions tend to equilibrate faster than metabolic reactions [52]. As a result, the distribution of species vary independently from the extension of metabolic reactions. Therefore, we simplify our analysis by lumping all the species together, representing them as a single reactant with respect to which mass-balance equations are expressed

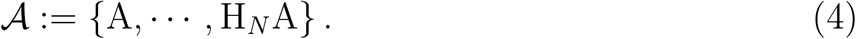

Here, 𝒜 represent a reactant in the system the concentration of which is given by

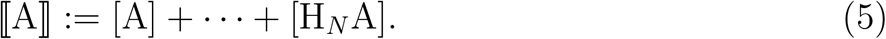

In subsequent sections, we sometimes denote the concentration of 𝒜 by *C*_𝒜_ to simplify the notation. For reactant 𝒜 with only one species A, ⟦A⟧ and [A] are indistinguishable.

For each species A′ ∈ 𝒜, the corresponding model fraction *ρ*_A′_ := [A′]*/* ⟦A⟧ is calculated from hydrogen-dissociation constants irrespective of the reactions A′ participates in [53].

As previously stated, the protocell membrane in our model is made of semi-permeable minerals to allow the exchange of materials between the protocell and its environment. The positive membrane potential developed across the membrane due to a pH gradient affects the transport rate of all charged species. The transport rate of charged or uncharged species through such membranes can be determined by solving the species mass-balance equation and Maxwell’s first law [1, 54]. We approximate the diffusive transport rates by [54]

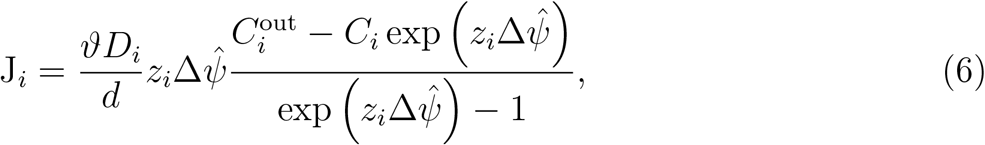

where 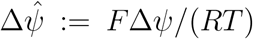 with Δ*ψ* the membrane potential, *ϑ* tortuosity coefficient, *F* Faraday constant, *T* temperature, and *R* the universal gas constant. Moreover, J_*i*_ denotes the diffusive flux, *D*_*i*_ the diffusivity coefficient, *z*_*i*_ the valence, *C*_*i*_ intracellular concentration, and 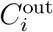 extracellular concentration of species *i*. Note that Eq. (6) is an approximate expression derived for a flat membrane, where the electric potential field *ψ* varies linearly along its thickness [54]. We may derive a more general expression for the membrane transport rate of reactant 𝒜 from Eq. (6) that accounts for other factors that contribute to its import flux

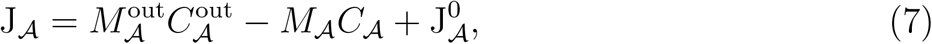

where

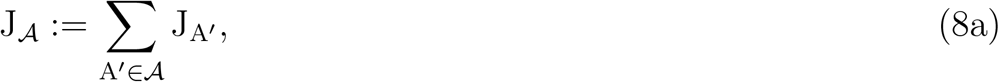

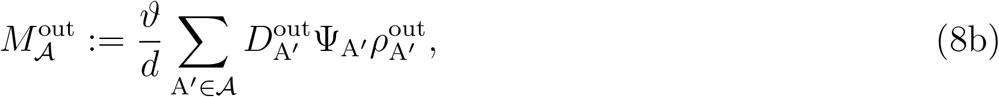

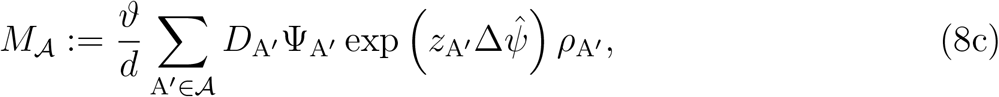

with 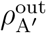 the mole fraction of A′ in the extracellular environment, 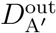 diffusivity coefficient of A′ in the extracellular environment, *ρ*_A′_ the mole fraction of A′ in the intracellular environment, *D*_A′_ diffusivity coefficient of A′ in the intracellular environment, and

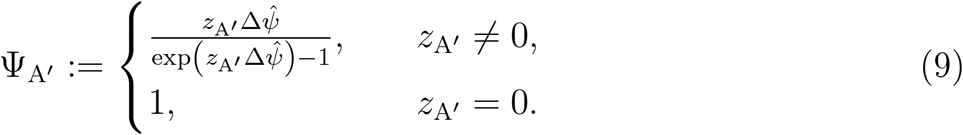

The constant term 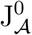 in Eq. (7) is a given parameter of the model. For all reactants except Fe^2+^, 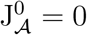. However, for Fe^2+^, 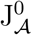 is the rate at which the iron-sulfide membrane dissolves into the intracellular environment (see Fig. S1C).

Assuming that all the species occupy a spherical region as they move, we approximate their diffusivity coefficient by Stokes–Einstein equation

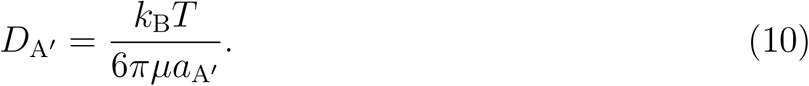

Here, *k*_B_ is the Boltzmann constant, *μ* the viscosity of water in the membrane, and *a*_A′_ the radius of the sphere that characterize A′. We used the Van der Waals radius of A′ to estimate *a*_A′_. Van der Waals radii were computed with Calculator Plugins, Marvin 20.21.0, 2020, ChemAxon (http://www.chemaxon.com). The Van der Waals radii of the unspecified thiol HS–R and its thioesters derivatives R^*/*^–COS–R were approximated assuming that the hypothetical substituent R is the alkyl group of ethanethiol.

Having introduced the definition of diffusive flux and concentration for species and reactants, we now can proceed to the formulation of mass-balance equations. Here, the idea is to express these equations with respect to reactants instead of species, solving them for reactant concentrations. A balance between reaction and membrane transport rates determines these concentrations. Reactions in our protometabolic network are divided into two groups: (i) Membrane reactions. The reductive carboxylation steps and the reaction for the regeneration of FD_rd_ are in this group (see “Model Description”). These reactions occur inside the protocell membrane. (ii) Bulk reactions. All reactions of the protometabolic network besides the membrane reactions belong to this group. These reactions occur inside the protocell. Accounting for membrane transport and both groups of reactions, the mass-balance equations for all the reactants involved in the protometabolic network are written

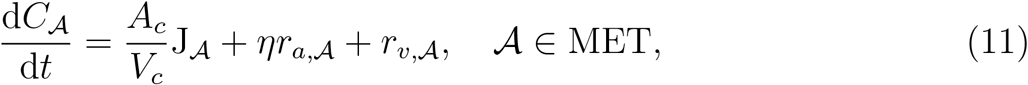

where *η* := 𝒱*/V*_*c*_ with *A*_*c*_ the inner surface area of the membrane, *V*_*c*_ volume of the protocell, 𝒱 volume of the membrane, *r*_*a*,𝒜_ production rate of 𝒜 by the membrane reactions, *r*_*v*,𝒜_ production rate of 𝒜 by bulk reactions, *t* time, and MET the index set of reactants that participate in metabolic reactions. Using the diffusive transport rates given by Eq. (7), Maxwell’s first law is implicitly applied in Eq. (11). Accordingly, the solutions of Eq. (11) simultaneously satisfy mass-balance constraints and Maxwell’s first law.

### Thermodynamic Constraints

Thermodynamic laws play a fundamental role in constraint-based models of metabolism [53, 55]. They dictate the limiting behavior of chemical reactions as they approach their equilibrium. The rate of reactions that are not in equilibrium are affected by their thermodynamic driving force [56]. As will be discuss in subsequent sections (see “Kinetic Constraints”), thermodynamic driving forces are determined by the equilibrium constants of chemical reactions.

Nonenzymatic reactions in the protometabolic network occur on catalytic surfaces similarly to enzymatic reactions of biochemistry. Mineral surfaces that catalyze these reactions in our model are made of transitions metals, which are positively charged in most cases [2]. The energetics of these reactions are affected by pH due to the interactions of hydrogen ions, protonation states of reactants, and charged surfaces. To determine the spontaneity of these reaction types, their Gibbs energy must be evaluated at a fixed pH [52]. This constraint can be incorporated into the Gibbs energy of reaction using the Legendre transformation. The quantity that results from the Legendre transformation of the Gibbs energy of reaction is referred to as the transformed Gibbs energy of reaction [52]. Its sign determines the spontaneity reactions for systems, where the temperature, pressure, and pH are the control parameters. As with the Gibbs energy of reaction, the transformed Gibbs energy of reaction at a given condition is evaluated with respect to a standard condition according to

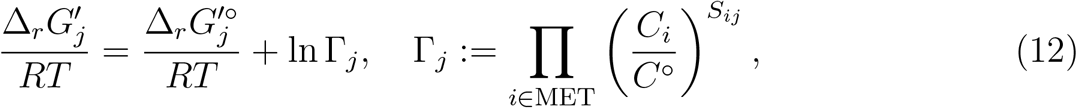

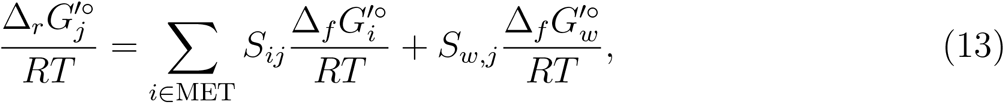

where *C*° is a reference concentration, *S*_*ij*_ are the components of the stoichiometry matrix of the protometabolic network, *S*_*w,j*_ stoichiometric coefficient of water in reaction *j*, 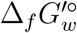 standard transformed Gibbs energy of formation of water, and 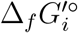 standard transformed Gibbs energy of formation of reactant *i* with 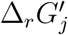 the transformed Gibbs energy, 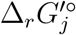 standard transformed Gibbs energy, and Γ_*j*_ quotient of reaction *j*. Note that, in this formulation, the concentrations of hydrogen ions and water do not affect the reaction quotients. Their stoichiometric coefficients do not contribute to *S* either. The standard transformed Gibbs energy of formation of a reactant 𝒜 is given by [53]

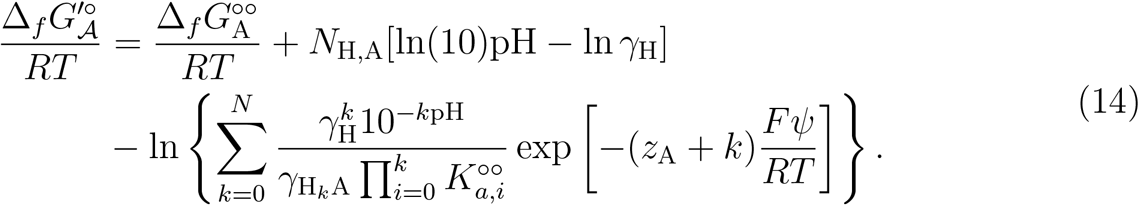

Here, *N*_H,A_ denotes the hydrogen content of the minimum charge state, 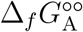 reference Gibbs energy of formation of the minimum charge state, *γ* activity coefficient of the species indicated by the subscript, 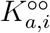 reference hydrogen-dissociation constant of species *i*, and *ψ* electric potential at which 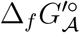 is evaluated. Note that, for each species *i*, the activity coefficient is a function of the ionic strength *I* of the solution it is in and its valence *z*_*i*_ [53]. Accordingly, the apparent equilibrium constant of reaction *j* is defined as

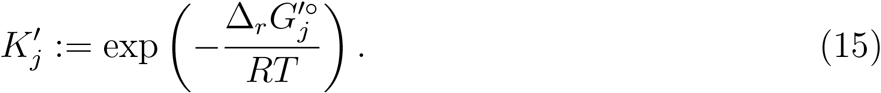

The activity coefficients in Eq. (13) depend on the ionic strength of the solution *I* and are approximated by the extended Debye–Hückel as described elsewhere [52, 53]. Finally, the thermodynamic constraints of the protocell model is expressed as

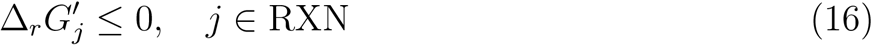

with RXN the index set of reactions in the protometabolic network.

From Eqs. (13)–(15), it is evident that 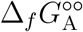 and 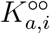 are required for all reactants to ascertain 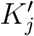. The value of these parameters for all the reactants of the protometabolic network is provided in Supplementary Data 1. We computed the reference Gibbs energy of formation of the minimum charge states using eQuilibrator (version 3.0) [57] and the reference hydrogen-dissociation constants using Calculator Plugins, Marvin (version 20.21.0, 2020), ChemAxon (http://www.chemaxon.com). In our protometabolic network, there are two nonstandard reactants for which 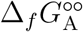 cannot be estimated from existing databases of tuned group-contribution parameters generated for standard metabolites of enzymatic metabolic pathways, namely hydroxyketoglutarate and oxalohydroxyglutarate. For these reactants, we estimated the reference Gibbs energy of formation of the minimum charge state using dGPredictor [58]. Both eQuilibrator and dGPredictor are web applications that estimate the standard transformed Gibbs energy of formation using group-contribution methods.

The energetics of the membrane reactions (*i*.*e*., those involving protoferredoxins; see “Mass Balance Constraints and Maxwell’s first law”) depend on the standard transformed Gibbs energy of formation of FD_rd_ and FD_ox_. In fact, the absolute values of these formation energies are not required. Providing the difference between these formation energies suffices to determine the standard transformed Gibbs energy of the membrane reactions. This difference may be expressed as

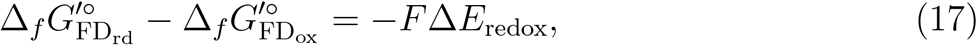

where Δ*E*_redox_ is the redox potential of protoferredoxins, which is a given parameter of the model. Once Δ*E*_redox_ has been provided, the standard transformed Gibbs energy of the membrane reactions can be specified.

Besides hydrogen ions, early metabolic reactions could also have been affected by metal-ion bindings. The early ocean is believed to have contained higher concentrations of several monovalent and polyvalent metal ions than modern seawater [59]. In particular, it would have contained appreciable amounts of ferrous iron in the absence of oxygen [17], which in turn could have interacted with other reactants that participated in early metabolic networks. These interactions would have altered the formation energy of reactants and the energetics of metabolic reactions. In this work, we only account for the binding of ferrous iron, as represented by

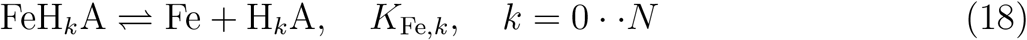

with *K*_Fe,*k*_ ferrous-iron binding constants. Note that, in this equation, the charge of compounds has not been shown for brevity. Accounting for the charge states arising from ferrous-iron biding reactions, we arrive at

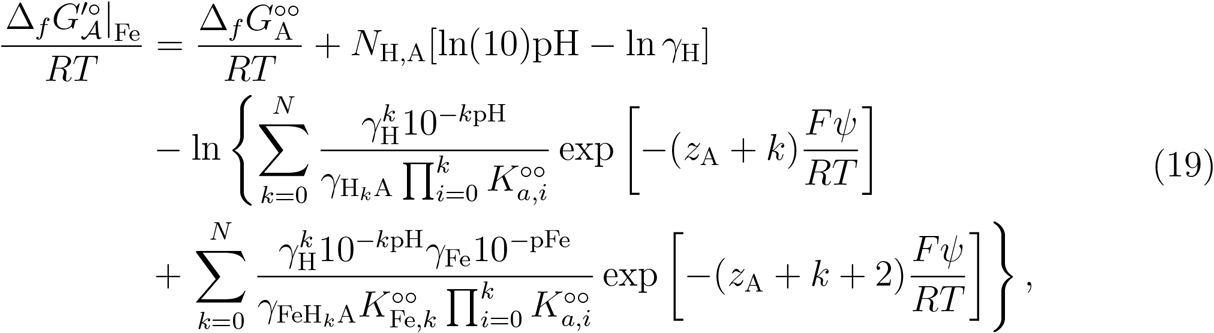

where 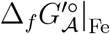 denotes the standard transformed Gibbs energy of formation of 𝒜 when the binding reactions of Fe^2+^ are accounted for. We use the notations Δ,_*r*_*G* ′| _Fe_ and *K*′|_Fe_ in a similar manner. Note that, in the main article, we dropped the subscript Fe when referring to Δ,_*r*_*G* ′| _Fe_. Accordingly, the transform Gibbs energy of reactions, which are shown in Figs. 2E and 3C, are based on the expression given by Eq. (19).

### Kinetic Constraints

All the reactions of the protometabolic network are surface-catalyzed. Therefore, to impose kinetic constraints, we require rate laws that account for reactant-catalyst bindings and surface reaction rates. We adopt Michaelis–Menten type expressions to quantify the kinetics of surface metabolic reactions in our model. There are eight reaction types in the protometabolic network: (i) thioesterification, (ii) membrane reactions (reductive carboxylation and protoferredoxin regeneration), (iii) retro-aldol and aldol, (iv) oxidative decarboxylation, (v) hydration and dehydration, (vi) reduction, (vii) decarboxylation, and (viii) solid catalyst synthesis. In this section, we present the rate laws that we used in our model for each reaction type to solve mass-balance equations Eq. (11). However, we first consider the Michaelis–Menten reaction mechanism

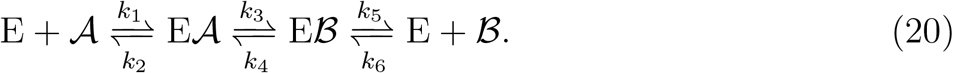

We use the general rate law for this reaction system to derive the rate laws for each reaction types in our model. Here, 𝒜, ℬ, and E represent the substrate, product, and catalyst of a given reaction in the network with *k*_*i*_ the rate constants of the individual steps. The overall rate law for this reaction system reads [56]

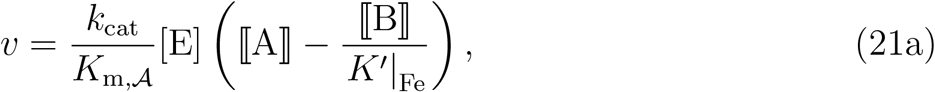

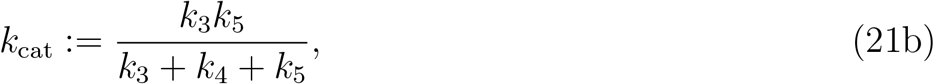

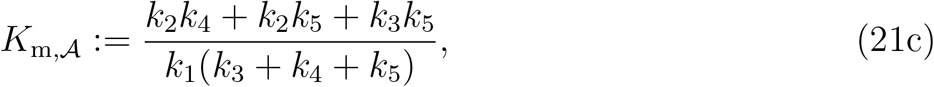

where [E] is the concentration of unbound catalyst, *k*_cat_ the turnover number, and *K*_m,𝒜_ is the Michaelis constant of 𝒜. As mentioned in the previous section, ferrous-iron binding constants are required to determine *K*′|Fe. Since these binding constants are generally unknown for most reactants in our protometabolic network, we simplify Eq. (19) in a way that allows us to parametrize them. To this end, we first write 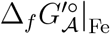 and 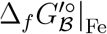 in terms of 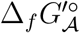 and 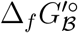, respectively. Let *T*_H,*A*_ and *T*_Fe,*A*_ stand for the first and second sum in the argument of the logarithm function in Eq. (19) for 𝒱. Let also Let *T*_H,*B*_ and *T*_Fe,*B*_ the same sums for ℬ. Then, we obtain the following standard transformed Gibbs energy of formation

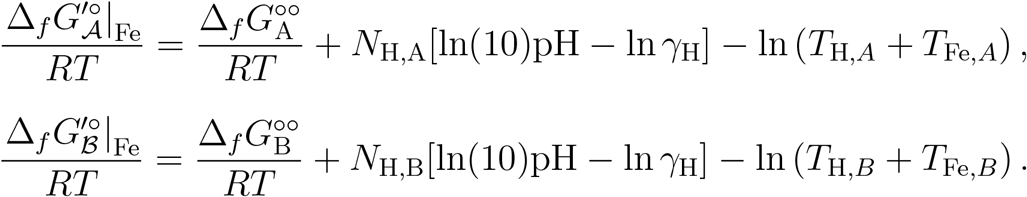

Assuming

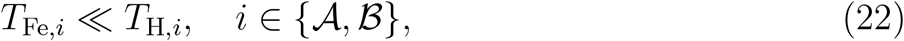

we have

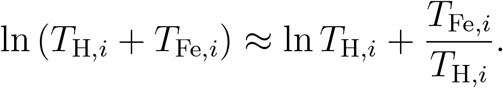

This assumption is justified by comparing the values of hydrogen-dissociation and metal-ion binding constants. Although we do not have all the binding constants for ferrous iron, we may estimate them from the binding constants of similar metal ions for which experimental data are available. The binding constants of similar divalent ions, such as Zn^2+^ and Mg^2+^, to a wide range of metabolites participating in biochemical reactions have been measured, and they are generally several orders of magnitude larger than hydrogen-dissociation constants of the same metabolites [60]. Thus, we expect that the binding constants of ferrous iron exhibit the same behavior for the reactants of the protometabolic network. Using the foregoing assumption, we obtain

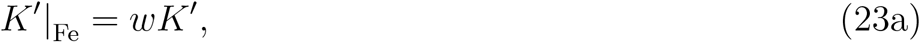

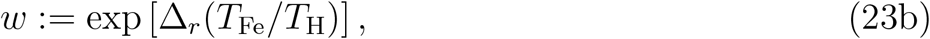

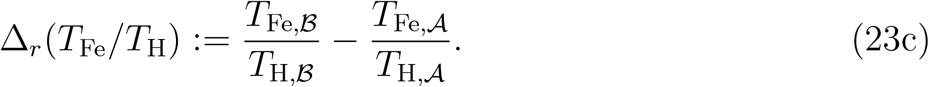

Substituting Eq. (23a) in Eq. (21a) yields

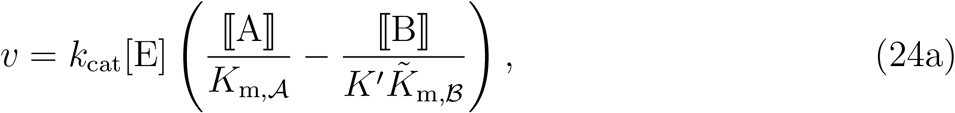

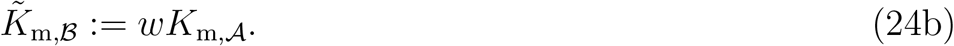

Here, *w* ∼ 𝒪(1) because |Δ_*r*_(*T*_Fe_*/T*_H_)| ≪ 1 according to the assumption stated in Eq. (22). Therefore, 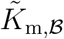 is of the same order of magnitude as *K*_m,𝒜_. Note that 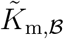 is not the same as the Michaelis constant *K*_m,ℬ_ that results from the reaction system Eq. (20). Thus, to avoid confusion, we refer to 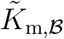 as the apparent Michaelis constant of ℬ at this point. However, as will be explained shortly, distinguishing between Michaelis and apparent Michaelis constants will not be necessary in the remainder of this document.

For a given reactant 𝒳, *K*_m,𝒳_ and 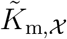 generally depend on the kinetic and Fe^2+^-binding constants of the reaction that 𝒪 participates in. However, to minimize the number of model parameters, we assume that the Michaelis and apparent Michaelis constants of a given reactant in the protometabolic network is the same for all reactions. Because of this assumption, we no longer need to distinguish between Michaelis and apparent Michaelis constants in the rate laws that will be discussed later in this section. Accordingly, we rewrite Eq. (24a) as

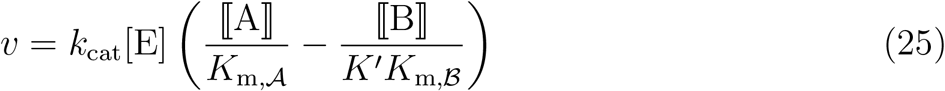

with *K*_m,𝒜_ and *K*_m,ℬ_ the Michaelis constant of the substrate and product, which are regarded as model parameters. In the following, we drive rate laws for each reaction type using Eq. (25). Note that reaction rates that are ascertained from ⟦A⟧ and ⟦B⟧ according to this rate law automatically satisfy the thermodynamic constraints, irrespective of the flux direction.

#### 1. Thioesterification

As previously stated, thioesterification reactions are catalyzed by H^+^ and Fe^2+^ in our model. Accordingly, the general reaction equation for thioesterification steps can be written

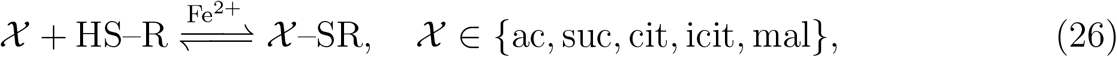

where 𝒳 represent a carboxylic acids and 𝒳–SR the respective thioester. Reactant identifiers are given in Supplementary Data 1. Following the development of Alberty [52], we do not explicitly state the stoichiometric coefficients of H^+^ and H_2_O that are consumed or produced in reaction equations—a convention we adhere to in the remainder this document. Given that all reactions occur in a dilute aqueous environment, the concentration of water can be regarded as a constant. Therefore, it only affects the reaction rates indirectly through their thermodynamic driving forces, or apparent equilibrium constants to be more specific (see Eq. (13)). Similarly, we assume that H^+^ only affects the reaction rates by altering the thermodynamic driving forces. The contribution of H^+^ is captured in Eq. (25) by using the apparent equilibrium constant to evaluate the thermodynamic driving force. The apparent equilibrium constant in turn accounts for the energetics of H^+^ being exchanged between reactants and the aqueous environment. Taking all these considerations into account, the rate of thioesterification reactions can be expressed as

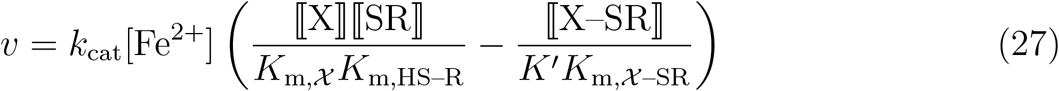

with X, SR, and X–SR the minimum charge states of 𝒳, HS–R, and 𝒳–SR, respectively. Recall, we adopted the notation ⟦minimum charge state⟧ to represent the concentration of a reactant (see “Mass Balance Constraints and Maxwell’s first law”).

#### 2. Membrane reactions

The reductive carboxylation steps of the protometabolic network occur inside the membrane on its acidic side. They are driven by protoferredoxin redox couples according to

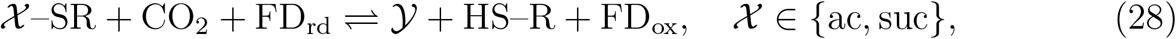

where 𝒳–SR is the thioester derived from the carboxylic acid 𝒳 𝒴 and the carboxylic acid resulting from the carboxylation of 𝒳. These reactions consume H^+^, so they are thermodynamically favorable in acidic solutions. Decreasing the intracellular pH increases their apparent equilibrium constants and their rates as a result. Moreover, protoferredoxins serve as both catalyst and reducing/oxidizing agents for these reactions. We use the rate law

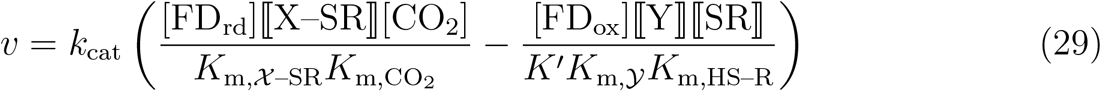

to describe the kinetics of reductive carboxylation reactions. Here, Y is the minimum charge state of 𝒴 with [FD_rd_] and [FD_ox_] the concentrations of FD_rd_ and FD_ox_, respectively. These concentrations are defined as the number of moles of the respective protoferredoxin unit cell per unit volume of the membrane. In this equation, we assume that the rate of forward and reverse reactions are proportional to the amount of FD_rd_ and FD_ox_ that are present in the membrane.

The reduced protoferredoxin FD_rd_, which is consumed by the reductive carboxylation reactions, is regenerated inside the membrane on its alkaline side

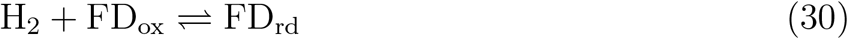

using H_2_ as the reducing agent. This reaction produces H^+^, and it is thermodynamically favorable in alkaline environments. The rate law for this reaction reads

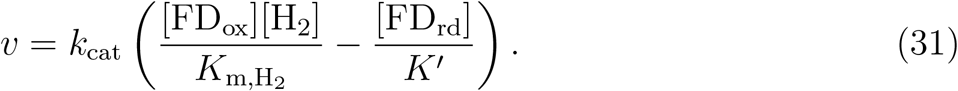

#### 3. Retro-aldol and aldol

Aldol and retro-aldol reactions play a significant role in biochemistry. They can be catalyzed by a wide range of organic and inorganic catalysts [61]. They can also be promoted by acids or bases depending on the tendency of the participating compounds to accept or donate H^+^ [62]. Among inorganic catalysts, transition metals [63] and chiral metal complexes [62] are of particular interest in the context of life’s metabolic origins. With regards to the rTCA cycle, FeS was shown to promote the retro-aldol cleavage of citrate into acetate and oxaloacetate [37]. Similarly, the Fe^2+^/Fe^0^ system was observed to catalyze several aldol reactions involving the intermediates of the rTCA cycle [10]. In our model, aldol and retro-aldol reactions are catalyzed by both FeS and Fe^0^, which are synthesized through the interaction of Fe^2+^ and HS^−^ in the protocell. Aldol reactions in the protometabolic network can be described by the general equation

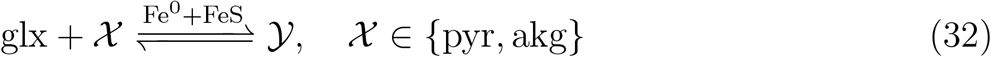

with 𝒳 a carboxylic acid and 𝒴 the carboxylic acid that results from the aldol condensation of glyoxylate and 𝒳. The rate law for these reactions is

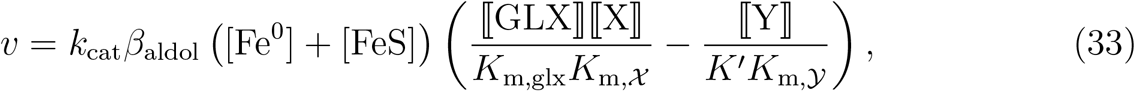

where GLX is the minimum charge state of glyoxylate. Retro-aldol reactions are described by

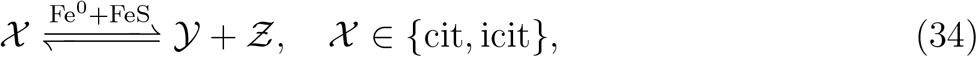

where 𝒳 is a carboxylic acid with 𝒴 and *Ƶ* the carboxylic acids that results from the retro-aldol cleavage of 𝒳. The rate law for these reactions read

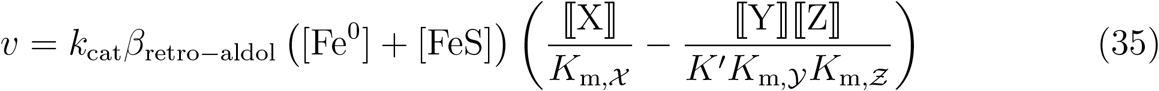

with Z the minimum charge state of *Ƶ*. In these rate laws, *β*_aldol_ and *β*_retro−aldol_ are molefractions of FeS-Fe^0^ assemblages that catalyze aldol and retro-aldol reactions, respectively.

#### 4. Oxidative decarboxylation

Oxidative decarboxylation steps are one group of off-cycle reactions that we accounted for in the protometabolic network. They are parasitic reactions since they counteract the reductive carboxylation steps, diminishing the efficiency of the carbon fixation. The competition between oxidative decarboxylation and reductive carboxylation reactions an important factor, determining whether a complete-cycle flux can be achieved. Previous studies on nonenzymatic analogues the TCA cycle detected these reactions in the presence of Fe^2+^ [10]. In the protocell model, these reactions are catalyzed by Fe^2+^ as indicated by

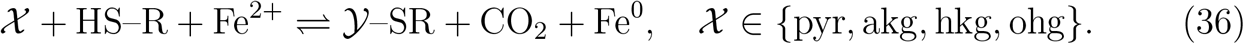

Here, 𝒳 represent a carboxylic acid and 𝒴 the carboxylic acid that results from the decarboxylation of 𝒳. Note that Fe^2+^ and Fe^0^ simultaneously serve as reducing/oxidizing agents and catalysts in these equations. The following rate law describes the kinetics of these reactions

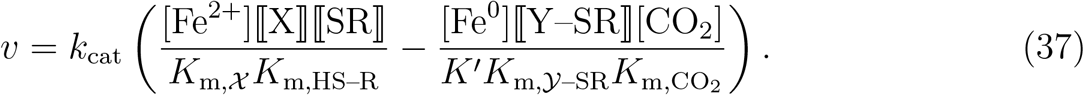

#### 5. Hydration and dehydration

Previous experimental studies showed that the hydration and dehydration steps of the rTCA and TCA cycles can be promoted by Fe^2+^, Zn^2+^, and Cr^3+^ without enzymes [9, 10]. Specifically, both Fe^2+^ and Zn^2+^ were observed to promote the dehydration of malate and isocitrate to fumarate and aconitate, respectively [9, 10]. However, the hydration of aconitate to citrate was observed only in the presence of Cr^3+^ [9]. In the protocell model, we consider the dehydration reactions

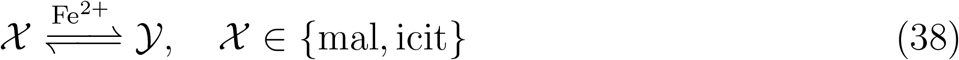

with the rate law

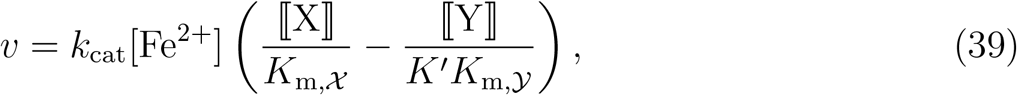

Where 𝒳 is a carboxylic acid and 𝒴 the product of its dehydration. The only hydration reaction in the protometabolic network is

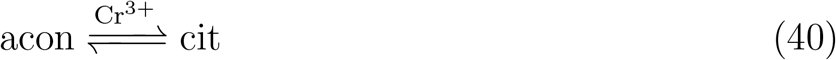

the rate law of which is given by

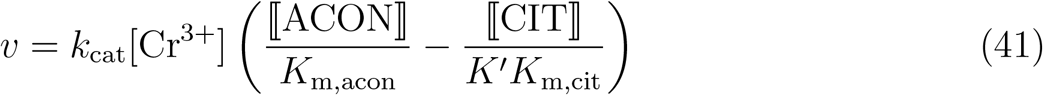

with ACON and CIT the minimum charge states of acon and cit, respectively.

#### 6. Reduction

In the absence of oxygen, the early Earth’s crust is believed to have been made of reducing mineral assemblages, mostly containing silicate, iron, nickle, and magnesium [17, 64]. Among these minerals, zero valent iron Fe^0^ and iron monosulfides FeS have been extensively studied in the origins-of-life literature as the main reducing agent [17, 65]. Specifically, Fe^0^ were shown to promote the reduction steps of the rTCA cycle [9]. In the protometabolic network, the reduction steps are also driven by Fe^0^ according as described by the following equation

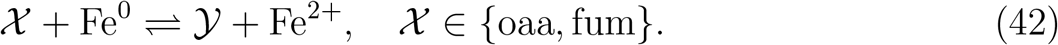

Here, 𝒳 is a carboxylic acid and 𝒴 the product of its reduction. The rate law for these reactions is

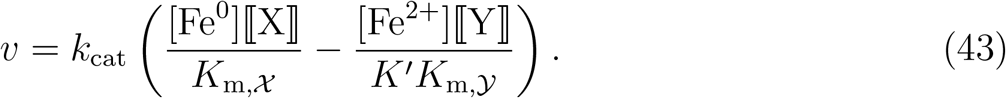

In these reactions, Fe^0^ and Fe^2+^ are both catalysts and reducing/oxidizing agents.

#### 7. Decarboxylation

Neutral decarboxylation steps are another group of off-cycle reactions that we considered in the protometabolic network. Nonenzymatic decarboxylation reactions in the presence of Fe^2+^ were observed previously in the prebiotic chemistry literature [10]. According, the only neutral decarboxylation step of the protometabolic network is catalyzed by Fe^2+^ as

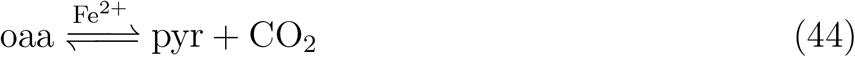

the rate law of which is

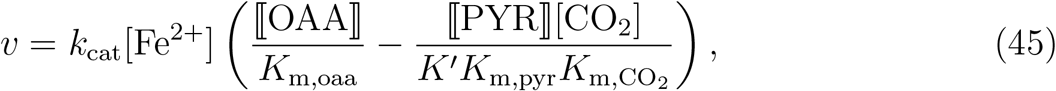

where OAA and PYR are the minimum charge states of oaa and pyr, respectively.

#### 8. Solid catalyst synthesis

In the discussion of the rate laws so far, we noted that Fe^0^ and FeS play the role of catalysts and reducing agents in the nonenzymatic reactions of the protometabolic network. As discussed in previous sections (see “Model Description”), FeS-Fe^0^ assemblages that precipitate on lime minerals inside the protocell as a result of the interaction of Fe^2+^ and HS^−^ are the sources of Fe^0^ and FeS that promote the foregoing protometabolic reactions. In the protocell model, assemblages of FeS-Fe^0^ in turn are synthesized from Fe^2+^ through

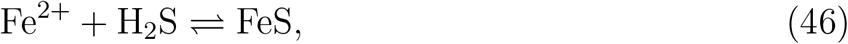

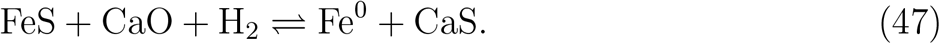

Reactions of the type in Eq. (47) were previously examined in the context of extractive metallurgy [66]. The rate laws for these reactions are given by

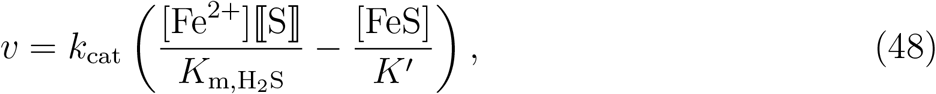

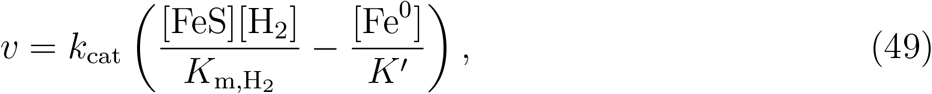

where S in Eq. (48) is the minimum charge state of H_2_S. Note that, CaO and CaS are produced and consumed only in Eq. (47), so [CaO] + [CaS] remains constant at all times. Therefore, to simplify our analysis, we do not solve their mass-balance equations to determine their concentrations. Since we are only interested in the solution of mass-balance equations near steady states in this work, we assume that CaO and CaS are abundantly available in the protocell, so their concentrations do not deviate form steady-state values significantly, and they approximately remain constant in time. Accordingly, [CaO] and [CaS] can be factored out and implicitly accounted for in *k*_cat_ and 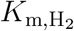 in Eq. (49).

Having detailed the kinetic constraints of the protocell model, we conclude this section by emphasizing two points, namely (i) inorganic reagents driving the protometabolic network and (ii) parametrization of the rate laws. With regards to the first point, all the steps of the protometabolic network are promoted by purely inorganic reagents. Catalysts, reducing/oxidizing agents, and energy sources are all either solid transition metals or inorganic ions in aqueous phase. Some of the regents serve multiple purpose. For example, Fe^0^ is a catalysts in some reactions and a reducing agents in others. Table S2 summarizes the characteristics of the inorganic reagents in the protocell model. Regarding the second point, in our formulation, the turnover number *k*_cat_ and Michaelis constant *K*_m_ are the key parameters that characterize the kinetics of nonenzymatic reactions. As will be explained in the next section, we determine the Michaelis constants by solving an optimization problem, which ensures that the mass-balance equations have a steadystate solution. However, we treat the turnover numbers as free parameters with respect to which the feasibility of first carbon-fixing cycles is to be investigated. Turnover numbers in turn measure the efficiency of catalysts that promote metabolic reactions. Therefore, we adopt this formulation to better understand the role of turnover numbers in the evolution of enzymes and genetic codes. The idea is to identify ranges turnover numbers that furnish stable, self-sustaining carbon-fixing cycles using parameter-sweep computations.

**Table S2:** Inorganic reagents, promoting the nonenzymatic reactions of the protometabolic network. The concentration of these reagents are either the variables of the model, which are determined from the mass-balance equations, or fixed parameters of the model.

| Reagent | Reducing | Oxidizing | Energy source | Catalyst | Type |
| --- | --- | --- | --- | --- | --- |
| $\Delta\text{pH}$ | ✗ | ✗ | ✓ | ✗ | Parameter |
| $\text{Cr}^{3+}$ | ✗ | ✗ | ✗ | ✓ | Parameter |
| $\text{H}^+$ | ✗ | ✓ | ✗ | ✓ | Parameter |
| $\text{H}_2$ | ✓ | ✗ | ✓ | ✗ | Variable |
| $\text{H}_2\text{S}$ | ✓ | ✗ | ✓ | ✗ | Variable |
| $\text{FeS}$ | ✓ | ✗ | ✓ | ✓ | Variable |
| $\text{Fe}^0$ | ✓ | ✗ | ✓ | ✓ | Variable |
| $\text{Fe}^{2+}$ | ✗ | ✓ | ✗ | ✓ | Variable |
| $\text{FD}_{\text{rd}}$ | ✓ | ✗ | ✓ | ✓ | Variable |
| $\text{FD}_{\text{ox}}$ | ✗ | ✓ | ✗ | ✓ | Variable |

Although we used the same symbol to denote the turnover number and Michaelis constants of every reaction, the values of these parameters are generally not the same for all reactions. As previously stated, for a given reactant 𝒜 in the protometabolic network, we assume that *K*_m,𝒜_ is the same for all the reactions that 𝒜 participates in. However, for turnover numbers, our goal is to examine the feasibility of the protometabolic network with respect to a single parameter *k*_cat_—the characteristic turnover number— that captures the order of magnitude of the turnover numbers for all the reactions in the network. The turnover number *k*_cat,*j*_ of a given reaction *j* in the network can deviate from the characteristic turnover number. However, the deviations

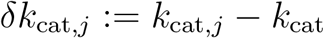

are assumed not to change the order of magnitude of *k*_cat,*j*_. That is, we assume that

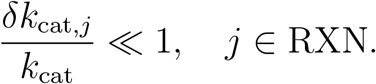

The underlying assumption of this approach is that the turnover numbers of the nonenzymatic reactions in the protometabolic network, which are all catalyzed by transition metals, are of the same order of magnitude. Although experimental data on the kinetic parameters of abiotic reactions in the context of life’s metabolic origins are scarce, this assumption is partly supported by a few kinetic studies on nonenzymatic analogues of core metabolic reactions. For example, Keller et al. [67] reproduced several steps of the TCA cycle, glyoxylate shunt, and succinic-semialdehyde pathway using a combination of iron sulfides and sulfate radicals, showing that the rate constants of the majority of the TCA reactions were in the same order of magnitude. In another work, Mayer et al. [14] studied the transamination reactions of glyoxylate with several *α*-keto acids using transition metals as catalysts and found turnover numbers of the order *k*_cat_ ≈ 10^−5^ 1*/*s for most cases. Moreover, the transition metals examined in this study enhanced the turnover number of glutamate transamination by a factor ≈ 10^2^ compared to the case where no catalyst was used.

Drawing on the results from the previous kinetic studies, we simplify our analysis by assuming that the turnover number of every reaction in the protometabolic network is the same as the characteristic turnover number *k*_cat_, neglecting all the turnover-number deviations (*δk*_cat,*j*_ ≈ 0, *j* ∈ RXN). Kinetic studies of transition-metal catalyzed counterparts of core metabolic reactions could provide more accurate estimates for *δk*_cat,*j*_ and benefit future systems-level kinetic models, such as that presented in this work. Such experimental efforts could complement theoretical approaches and help elucidate the role of catalytic efficiency, as measured by turnover numbers, in the emergence and evolution of first metabolic cycles.

To investigate the role of turnover numbers, we also require a reasonable range for the turnover numbers of abiotic metabolic reactions catalyzed by transition metals. Reasonable estimates for these turnover numbers can be obtained by comparing the efficiency of transition-metal catalyzed and enzyme-catalyzed reactions. Enzymes are highly optimized catalysts. On average, they can enhance the rate of uncatalyzed chemical transformations approximately by 10^12^–10^16^ fold and 10^7^–10^8^ fold at 25° C and 100° C, respectively [68]. In contrast, inorganic catalyst have comparatively poor efficiencies, and generally they are not expected to increase the turnover numbers of uncatalyzed reactions by more than 10^2^ fold [4]. Given the temperature of early AHVs (see Fig. 1A) and using the median turnover number *k*_cat_ ≈ 10 1/s of enzymatic reactions [69] as a basis for comparing inorganic and organic catalysts, we find the range *k*_cat_ ≈ 10^−5^–10^−4^ 1/s for the turnover number of nonenzymatic reactions in our model, which is the range in which we examined the feasibility of the protometabolic network (see Fig. 2A).

## Steady States of Protocell

Once the rates of reactions in the protometabolic network have been ascertained from the rate laws outlined in the previous section, the production rates of reactants in the network are determined from

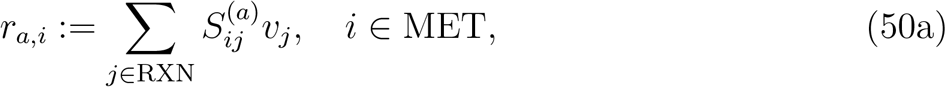

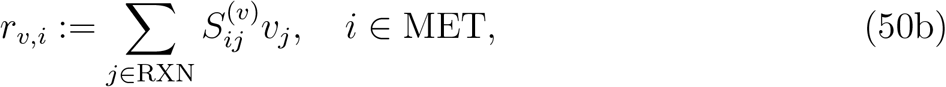

where 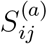 and 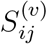 are the components of the stoichiometry matrices associated with the membrane and bulk reactions, respectively. Substituting Eq. (50) in Eq. (11), we obtain a closed system of ordinary differential equations from which the time-dependent concentrations of all the reactants in the protometabolic network are determined. This system has 27 variable concentrations and 24 metabolic fluxes. However, the system is not linearly independent when the rate laws outlined in the previous section are used in the mass-balance equations. The linear dependency arises because of the formulation of protoferredoxins in our model. Since the same amount of FD_rd_ is always consumed as FD_ox_ is produced or vice versa, the mass-balance equations for FD_rd_ and FD_ox_ are linearly dependent. To resolve this problem, we replace the mass-balance equation for FD_ox_ with a constraint on the total amount of protoferredoxins that the membrane can contain

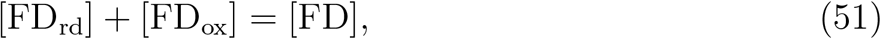

where

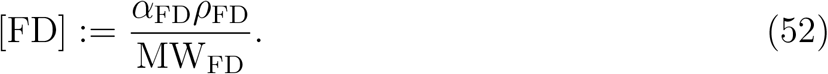

Here, *α*_FD_ the mass fraction of protoferredoxins in the membrane, which is a model parameter, with *ρ*_FD_ and MW_FD_ the density and molecular weight of protoferredoxins. Since the detailed structure and composition of protoferredoxins are unspecified in our model, we use the density and molecular weight of greigite to approximate *ρ*_FD_ and MW_FD_.

The main objective of this paper is to answer whether there are conditions under which the protometabolic network can reach steady states in the protocell outlined in Section “Model Description” (see also Fig. 1B), and if so, whether state states are stable. In particular, we seek ranges of physico-chemical parameters for which the mass-balance equations have steady-state solutions. The membrane potential Δ*ψ* and characteristic turnover number *k*_cat_ are the two parameters with respect to which we study the feasibility of the protometabolic network. Accordingly, our goal is to determine whether the system

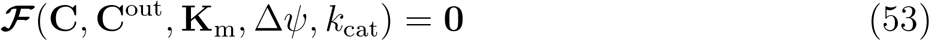

has a solution for any ranges of parameters in the (Δ*ψ, k*_cat_) parameter space. Here, ℱ is a vector, containing the right-hand-sides of mass-balance equations in Eq. (11), **C** the vector of concentrations, **C**^out^ the vector of extracellular concentrations, and **K**_m_ the vector of Michaelis constants. Since Eq. (53) is a nonlinear system of equations, we formulate a phase-I optimization problem to determine if it has a solution for any values of parameters [70, 71]. Our preliminary numerical experiments indicated that, regarding the extracellular concentrations of reactants, membrane potential, turnover numbers, and Michaelis constants as fixed parameters, the system Eq. (53) is highly constrained for ranges of these parameters that are pertinent to AHV conditions on the early Earth, and in most cases, solving the phase-I problem for Eq. (53) proved fruitless. Therefore, we sought to find extracellular concentrations and Michaelis constants within reasonable ranges, such that the fundamental constraints of the protocell model are not violated. This alternative viewpoint is motivated by the fact that the exact values of the extracellular concentrations and Michaelis constants of reactants involved in early carbon-fixing networks under the AHV conditions of the early Earth are unknown. Accordingly, we solve the optimization problem

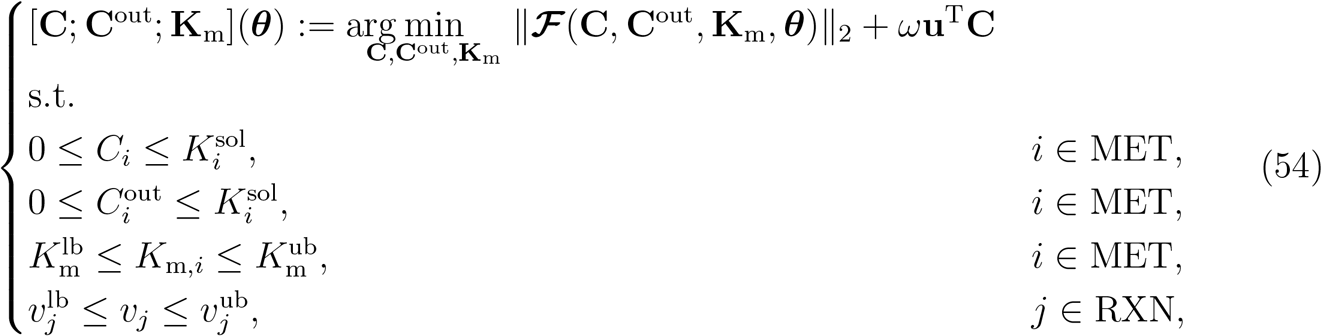

where ***θ*** := [Δ*ψ*; *k*_cat_] with 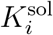 the solubility of reactant *i* in water, 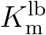 lower bound on Michaelis constants, 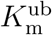 upper bound on Michaelis constants, 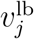 lower bound on the flux of reaction *j*, and 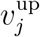 upper bound on the flux of reaction *j*. We approximate the lower and upper bounds on the Michaelis constants of the protometabolic network based on enzymatic Michaelis constants. Under physiological conditions, enzymatic Michaelis constants vary over a wide range, spanning several orders of magnitude from ∼ 10^−8^ M to ∼ 10^−1^ M [72]. Using these values as guide, we found steady-state solutions for Eq. (11) within reasonable physical range by choosing 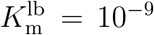 M and 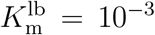. We also impose upper and lower bounds on some metabolic fluxes to eliminate inconsistent solutions that are not ruled out by the physico-chemical constraints of the protocell model. The only examples are the oxidative decarboxylation of pyruvate to acetate and of *α*-ketoglutarate to succinate. These steps like others in the protometabolic network are treated as reversible reactions in the protocell model. Although, in principle, they can proceed in the reverse direction without violating the thermodynamic constraints, the physico-chemical characteristics or crystal structure of the redox couple Fe^2+^/Fe^0^ that drive these reactions do not permit the reverse direction. Accordingly, for these reactions, we impose the lower bound 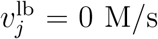 to ensure that the flux from acetate to pyruvate and from succinate to *α*-ketoglutarate only passes through the reductive carboxylation reactions (Fig. 1C, steps A and G), which are mediated by protoferredoxins, and not the reverse direction of the oxidative decarboxylation steps (Fig. 1C, steps V and X). We impose no upper or lower bound on the metabolic flux of other reactions (*i*.*e*., 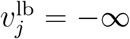 and 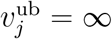). For these reactions, the concentrations are bounded, the rate laws automatically ensure that the metabolic fluxes are bounded.

Although Eq. (53) is a balanced system of equations, it could have multiple solutions for a given set of parameters because it is a nonlinear system. Therefore, we incorporate a regularizer into the optimization problem Eq. (54) that allows us to choose a unique solution based on a given criterion. In the objective function of Eq. (54), the first term measures deviations from the steady-state mass-balance constraints Eq. (53) and the second term is a linear regularizer, which penalizes deviations from the criterion based on which a unique solution is to be determined. Here, *ω* is the weight of the regularizer and **u** is a constant vector that specifies the criterion. For example, through numerical experiments, we found that choosing a regularizer that maximizes the concentration of *α*-ketoglutarate leads to steady-state solutions that correspond to a complete-cycle flux in most cases.

We determine the stability of steady-state solutions obtained from Eq. (54) by computing the eigenvalues of the Jacobian matrix 𝒥 defined as

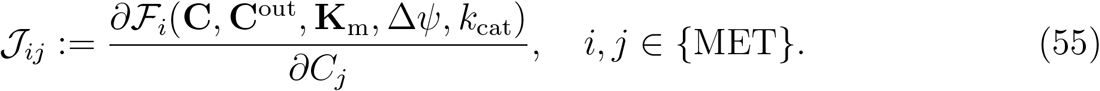

Note that, once we have found **C**^out^ and **K**_m_ that allow the protometabolic network to attain a steady state, we treat them as fixed parameters in the stability analysis. According to linear stability theory [73], a steady-state solution of Eq. (11) is linearly stable if the eigenvalues of the Jacobian matrix all have negative real parts. Otherwise, the steady-state solution is unstable.

A stable steady-state solution of Eq. (11) is an interior point of a stability region in the parameter space. To determine parts of the (Δ*ψ, k*_cat_) parameter space in which steady-state solutions are stable and those in which steady-state solutions are unstable, we use branch-continuation methods [73]. Accordingly, we compute steady-state solutions of Eq. (11) over a nonuniform grid in the parameter space by performing branch continuation along Δ*ψ*- and *k*_cat_-directions from a starting interior point provided by the optimization problem Eq. (54). Steady-state solution branches along Δ*ψ*- and *k*_cat_-directions are obtained by solving

**Table S3:** Model parameters.

| Parameter | Value | Parameter | Value |
| --- | --- | --- | --- |
| $T_{\text{out}}$ | 150° C | $R_c$ | $10^{-7}$ m |
| $T_{\text{in}}$ | 90° C | $d$ | $2 \times 10^{-8}$ m |
| $\text{pH}_{\text{out}}$ | 9 | $[\text{Cr}^{3+}]$ | $10^{-3}$ M |
| $\text{pH}_{\text{in}}$ | 5 | $\alpha_{\text{FD}}$ | 0.8 |
| $I_{\text{out}}$ | 0.3 M | $\beta_{\text{aldol}}$ | 0.8 |
| $I_{\text{in}}$ | 0.3 M | $\beta_{\text{retro-aldol}}$ | 0.1 |
| $J_{\text{Fe}^{2+}}^0$ | $10^{-2}$ mol/(m<sup>2</sup>s) | $\vartheta$ | $10^{-5}$ |
| $\Delta E_{\text{redox}}$ | 1 V | $C_t^{\text{out}}$ | 0.4 M |

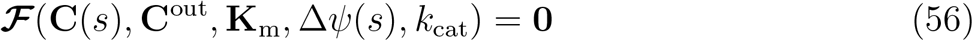

and

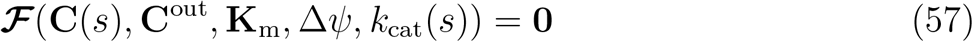

using arclength continuation methods, respectively. Here, *s* denotes the arclength along solution the respective branch. We refer the reader to the work of Seydel [73] for a comprehensive overview of arclength-continuation methods and how solution branches are constructed from Eqs. (56) and (57). Using these techniques, we constructed the stability region discussed in the main article (see Fig. 2A) and characterized the bifurcation of steady-state solution branches along its boundaries. We found that the stability region is bounded in the membrane potential and unbounded in the turnover number in the ranges of parameters that we studied. Specifically, solution branches along Δ*ψ*-directions bifurcate and lose stability at both small and large membrane potentials but not along *k*_cat_-directions. The stability-region boundary at small membrane potentials correspond to turning points and at large membrane potentials to simple branch points. The Jacobian matrix is singular at both boundaries, that is rank(ℱ_**C**_) = *n*−1 with 𝒥 := ℱ_**C**_ (subscript indicates the variables with respect to which derivative is taken; see Eq. (55)) and *n* := MET the number of variable concentrations. We can characterize the type of bifurcation along the stability-region boundaries based on the rank of the following concatenated matrix for solution branches that are constructed from Eq. (56)

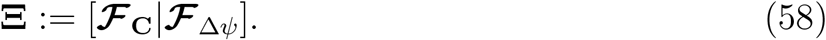

Accordingly, rank(**Ξ**) = *n* at turning points and rank(**Ξ**) = *n*−1 at simple branch points (see Definitions 2.7 and 2.8 of Seydel [73]). We can follow a similar approach for solution branches that are constructed from Eq. (57). We also characterized the eigenvalues of the Jacobian matrix throughout the stability region and found that dynamic trajectories around steady-state solutions can exhibit oscillatory or nonoscillatory behaviors at different parts of the stability region (Fig. S6). All the model parameters we used to perform all the computations in this paper are summarized in Table S3.

The solubility of reactants in Eq. (54) were computed with Calculator Plugins, Marvin (version 20.21.0, 2020), ChemAxon (http://www.chemaxon.com). We formulated the global optimization problem Eq. (54) in GAMS (version 28.2.0, 2019) and solved it using the global solver BARON (version 19.7.13, 2019) equipped with the local nonlinear programming solver CONOPT (version 3.17K, 2019) and the linear programming solver CPLEX (version 12.9.0, 2019). Branch-continuation problems Eqs. (56) and (57) were solved using custom codes in MATLAB (version R2019b). All the custom codes written in GAMS and MATLAB to generate the results presented in this paper are provided in Supplementary Code 1.

**Figure S6:**
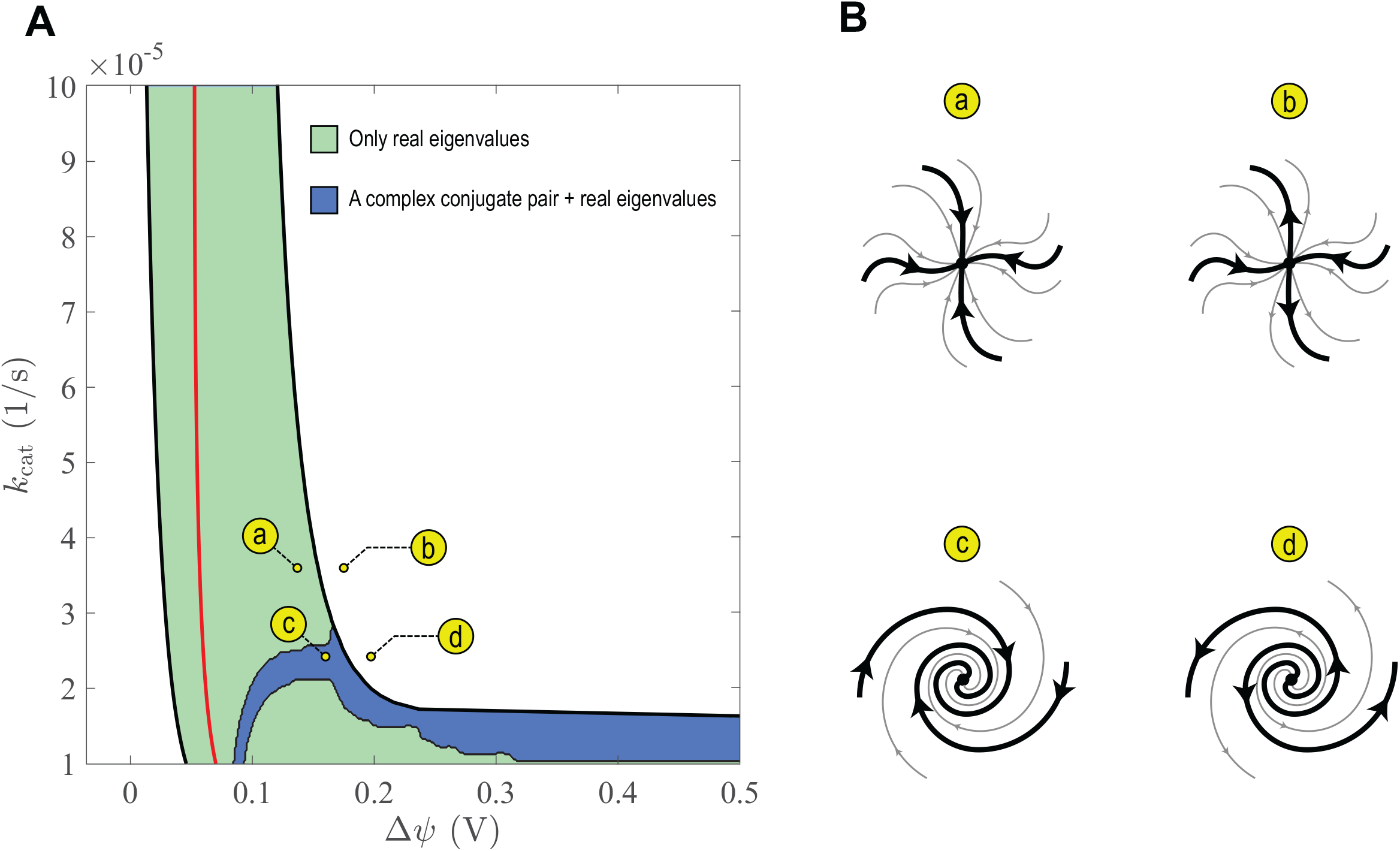
Near-steady-state dynamics of the protocell model. (A) Characteristics of the eigenvalues of the Jacobian matrix in Eq. (55) in the stability region of the protocell discussed in the main article (see Fig. 2A). (B) Schematic representation of dynamic trajectories near steady-state solutions in a two-dimensional projection of the phase space at selected points of the parameter space.

### Osmotic Pressure Drives Protocell Growth

As we stated in the main article, the membrane potential is the main driving force of growth in our model. The membrane potential is positive due to the direction of the pH gradient across the protocell membrane. Because the positive membrane potential nonuniformly distributes ions across the membrane, an osmotic pressure differential is developed in the protocell. Note that the osmotic pressure differential is generated because of the uneven movements of ions to one side of the membrane whether or not metabolic reactions occur inside the protocell. Accordingly, our goal in this section is to estimate the change in the osmotic pressure differential as a result of metabolic reactions.

We approximate the osmotic pressure of an aqueous mixture of solutes in water using a truncated Pitzer model [74]

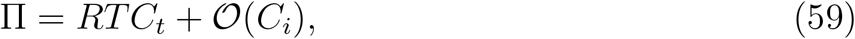

where Π is the osmotic pressure of the aqueous solution of interest, *C*_*t*_ total solute concentrations, and *C*_*i*_ passed to the order-of-magnitude operator 𝒪 is a representative solute concentration. Following the approach we introduced in our earlier work [1], we partition all solutes that contribute to the osmotic pressure into reactive and nonreactive solutes. Since Eq. (59) is linear in solute concentrations, we can decompose the total osmotic pressure into a component induced by reactive and another induced by nonreactive solutes and compute each component separately. Let 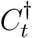 and 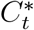 denote the total concentration of reactive and nonreactive solutes in the intracellular environment. Let also 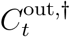 and 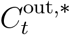 denote the same quantities in the extracellular environment. Accordingly, we can consider the following decompositions

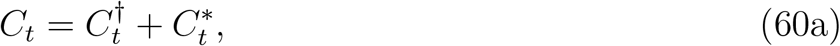

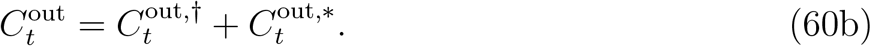

First, we estimate the osmotic pressure differential due to nonreactive species. To simplify the analysis, we assume that the contribution of reactive solutes to the osmotic pressure of the extracellular environment is negligible compared to that of nonreactive species, so that 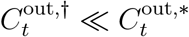 and 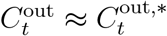. We also assume that the extracellular environment is electroneutral and contains equal amounts of negatively and positively charged solutes, which are all monovalent. Thus,

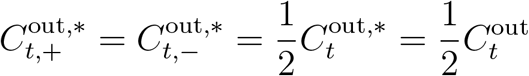

with 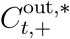 and 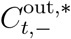 the total concentration of positively and negatively charged solutes in the extracellular environment. At steady state, the membrane transport rate of nonreactive species is zero [1]. Using this property and the expression for diffusive membrane transport rate given by Eq. (6), we can ascertain the total concentration of positively and negatively charged solutes in the intracellular environments

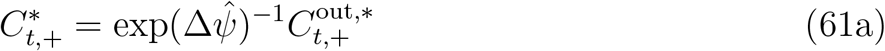

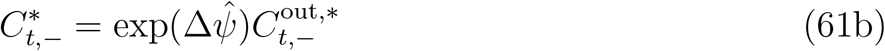

from which it follows that

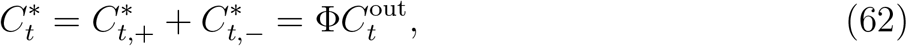

where

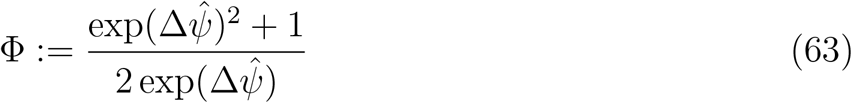

with 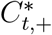 and 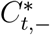 the total concentration of positively and negatively charged solutes in the intracellular environment. Therefore, the contribution of nonreactive solutes to the osmotic pressure differential can be obtain using Eq. (59)

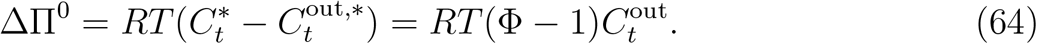

Next, we obtain the following expression for the osmotic pressure differential due to both reactive and nonreactive solutes using Eq. (59)

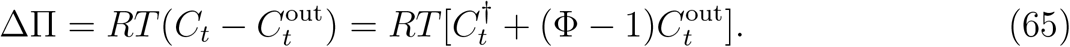

The iron-sulfide membrane in our protocell model is rigid. Therefore, an increase in the osmotic pressure differential due to the accumulation of protometabolic network products, as measured by ΔΠ*/*ΔΠ^0^, results in an excess in-plan stress throughout the membrane without changing the volume of the protocell. However, as we previously discussed (see “Model Description” and Fig. S2), at early stages of evolution, lipid membranes could have replaced iron-sulfide membranes once lipid-synthesis pathways had been incorporated into first carbon-fixing cycles. In these early cell-like structures with stretchable membranes, an increase in the osmotic pressure differential could have resulted in a volume expansion and been a driving force of growth. Therefore, it is useful to measure the ratio ΔΠ*/*ΔΠ^0^ in terms of the volume expansion that it would cause in a deformable cell as it is more relevant to selective processes underlying transitions from iron-sulfide to lipid membranes. To this end, we compare the protocell model, which has a rigid membrane, to a hypothetical protocell with a deformable membrane. Let 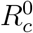 denote the radius of the protocell model with 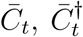 and 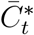 the total concentration of all solutes, reactive solutes, and nonreactive solutes in its intracellular environment. We then consider a hypothetical protocell with the same radius 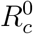 as the protocell model, only containing the nonreactive solutes of the protocell model. We assume that the membrane of the hypothetical protocell is elastic with a stretching rigidity *γ*. Regarding the total concentration of nonreactive solutes in the extracellular environment 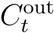 as a fixed parameter (recall, we neglected 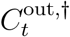 compared to 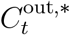, so the total concentration of nonreactive solutes and the total concentration of all solutes in the extracellular environment are the same, that is 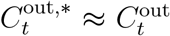), the total concentration of nonreactive solutes in the intracellular environment of the hypothetical protocell is the same as that of the protocell model, which is 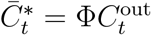 (see Eq. (62)). The goal here is to determine how much this hypothetical protocell would expand, if it contained the total amount of the reactive solutes of the protocell model 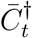.

To determine how much the volume of the hypothetical protocell expands as a result of the accumulation of the reactive solutes, we introduce two states: (i) The base state, which is the hypothetical protocell with an elastic membrane of stretching rigidity *γ* and radius 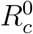. In this state, the total amount of solutes in the intracellular environment is 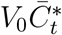, the osmotic pressure differential is denoted ΔΠ^0^, and 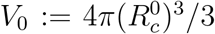. (ii) The expanded state, which is the hypothetical protocell with an elastic membrane of the same stretching rigidity as in the base state and radius 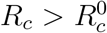, containing both the reactive and nonreactive solutes of the protocell model. In this case, the total amount of solutes in the intracellular environment is 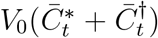 and the osmotic pressure differential is denoted ΔΠ. Therefore, the total concentration of solutes in the intracellular environment is 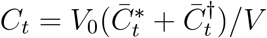 with 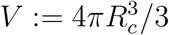, so we have

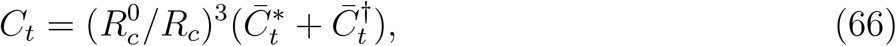

where *C*_*t*_ is the total concentration of solutes in the intracellular environment for the expanded state.

Assuming that the membrane in the hypothetical protocell is thin and elastic, the osmotic pressure differential is inversely proportional to the protocell radius [75]. Implementing this proportionality in Eqs. (64) and (65) yields

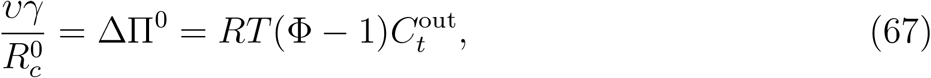

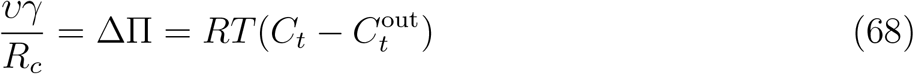

with *υ* the proportionality constant. Solving Eq. (67) for *γ* and substituting it in Eq. (68) along with *C*_*t*_ from Eq. (66) results in the cubic equation

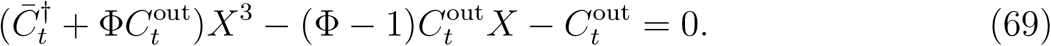

Here, 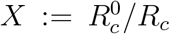 is the ratio of the radii of the base and expanded states, serving as a measure for volume expansion due to the accumulation of reactive solutes in the protocell. Note that 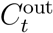 in this equation is a fixed parameter, Φ is a function of Δ*ψ*, and 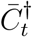 is determined from the steady-state solutions of the mass-balance equations Eq. (11). Therefore, for any point in the (Δ*ψ, k*_cat_) parameter space, there is a corresponding 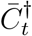 and *X*. Accordingly, as a specific example, we computed *X* throughout the stability region of the protocell model in the parameter space to gain general insights into the driving forces of growth for compartmentalized and self-sustaining carbon-fixing cycles at the origin of metabolism. The results are shown in Fig. 3A of the main article for 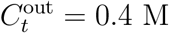.

